# Conditional feature importance with error control for machine-learning-based biomarker discovery in Alzheimer’s disease

**DOI:** 10.64898/2026.11.26.769462

**Authors:** Joseph Paillard, Erica Silvestri, Julien Bryois, Tony Kam-Thong, Gilbert Di Paolo, Angel Reyero Lobo, Ahmad Chamma, Erik Stomrud, Sebastian Palmqvist, Shorena Janelidze, Niklas Mattsson-Carlgren, Bertrand Thirion, Denis A. Engemann, Swedish BioFINDER study group and for the Alzheimer’s Disease Neuroimaging Initiative

## Abstract

Alzheimer’s disease (AD) is characterized by complex biology captured by proteomics, transcriptomics, genetics, and brain imaging. These high-dimensional and correlated data make biomarker discovery challenging. To disentangle biomedical AD data, we propose Conditional Feature Importance (CFI), a theory-grounded framework quantifying the unique contribution of individual or grouped variables in any machine learning model while controlling false discoveries. Across ADNI, BioFINDER, and UK Biobank (nearly 48,000 participants), CFI uncovered a striking contrast between unique versus redundant biomarkers across protein and imaging signatures of AD diagnosis, onset, and progression. Age and neurofilament light chain (NfL) were important for all three outcomes and A*β*_42_ was common to diagnosis and progression. Beyond these shared predictors, in BioFINDER, markers associated with neuronal and synaptic injury, including plasma total tau, CSF *α*-synuclein, and CSF neurogranin, contributed unique predictive information for diagnosis, whereas markers related to the amyloid cascade, CSF ptau181 and A*β*_40_ contributed unique predictive information for progression. *APOE ε*4 offered no unique information once downstream proteins were accounted for, whereas contextual markers such as CSF A*β*_40_ emerged only through interactions with other biomarkers. We argue that CFI increases the scientific value of machine learning by providing rigorous statistical insights on complex datasets.

## 1 Introduction

The etiology of Alzheimer disease (AD) is multifactorial and highly polygenic, involving more than a hundred associated genetic loci, interacting biological processes, and protein deregulation cascades [Uffelmann et al., 2026]. AD is a heterogeneous disorder; the effect size of prominent genetic risk factors is modulated by other patient characteristics, while frequent co-pathologies, including Lewy body and TDP-43 pathology, further contribute to diverse clinical manifestations [Young-Pearse et al., 2023; Knopman et al., 2021]. Combining diverse technologies, such as proteomics, neuroimaging, genetics, and demographic data, is necessary for a holistic understanding of the disease that is essential for developing better diagnostics and novel therapies, as each approach accesses complementary aspects of a shared underlying pathophysiology [Frisoni et al., 2025]. Furthermore, the clinical manifestation of AD is not strictly deterministic; modifiable lifestyle factors (e.g., education, hypertension, or type-2 diabetes) appear as epidemiological risk factors that may modulate the risk conferred by genetics and neuropathology [Livingston et al., 2024; Stern, 2012]. Consequently, the impact of any single characteristic can only be understood when evaluated within the participant’s broader biological and environmental context.

To better capture the complexity of AD and its progression, novel modeling approaches must be capable of capturing potentially complex interactions between variables. This emphasizes the importance of advanced statistical modeling approaches that can handle complex nonlinear relationships between many variables. The challenge is further compounded by the fact that multimodal biomedical datasets are inherently high-dimensional and contain highly correlated variables. This is, in particular, the case in a neurodegenerative condition like AD, where multiple types of brain imaging, cellular, and biofluid datasets are collected to characterize the disease. Consequently, it becomes difficult to discern the individual role of any single variable. Machine learning (ML) models excel at addressing these high-dimensional challenges and predicting biomedical outcomes from heterogeneous inputs. In particular, tabular foundation models, novel deep learning models pretrained to predict from heterogeneous data tables, have recently emerged as leading benchmarks on tabular data. They can learn complex representations and capture synergistic interactions, yielding more accurate predictions from heterogeneous biomedical inputs [Hollmann et al., 2025; Qu et al., 2025]. However, the adoption of such flexible models introduces a fundamental tension between predictive accuracy and scientific interpretation. Linear statistical models, commonly used in biomedical research, have limited capacity to uncover non-linear and interaction effects from large datasets. On the other hand, complex ML models capable of capturing complex interaction patterns come at the expense of transparency and statistical guarantees. Modern deep learning architectures feature millions of parameters to extract complex representations of the data, effectively rendering them opaque “black boxes”. To resolve this tension, we propose conditional feature importance (CFI) as a framework for estimating variable importance that quantifies the unique predictive information contributed by each variable, with formal guarantees against false discoveries. The CFI approach remains entirely agnostic to the chosen machine learning model and isolates the features that carry unique predictive information.

We will argue that this variable-importance framework is particularly well-suited for the AD field, which has transitioned from defining the disease solely as a clinical syndrome to a biological construct [Knopman et al., 2021]. The biological definition of AD relies on clinical imaging and selected fluid biomarkers, yet its refinement remains an active area of research. As new types of data are explored with ML, measuring variable importance accurately can translate into novel biomarker hypotheses. In this context, it is essential that importance is assigned only to variables that provide exclusive information about the underlying biology or pathology [Reyero-Lobo et al., 2025]. Yet, most standard feature importance methods fail this requirement in the presence of correlated variables, which are a hallmark of multimodal AD data. This can lead to misidentification of irrelevant variables as important predictors of disease (i.e., false discoveries) [Strobl et al., 2008; Chamma et al., 2023].

This shortcoming is rooted in how feature importance has been defined historically. Model interpretation has long relied on permutation-based measures and Shapley values imported from cooperative game theory [Breiman, 2001; Covert et al., 2020]. Such methods, however, remain heuristics: they were introduced to measure how much a given model relies on a feature rather than estimating a property of the data itself. Such methods, therefore, conflate marginal importance, the predictive power a feature carries on its own, with conditional importance, the predictive power a feature adds once all other measured features are available. CFI resolves this ambiguity by reframing the question as the estimation of a well-defined mathematical quantity, rooted in sensitivity analysis [Homma and Saltelli, 1996; Williamson et al., 2021], and backed by mathematically provable statistical guarantees. By construction, only features that bring information beyond the others receive positive conditional importance. Redundancy thus becomes a testable hypothesis, with CFI controlling the risk of false discoveries even when features are strongly correlated. CFI thereby turns powerful predictive models into instruments for scientific discovery.

In this work, we harness CFI to identify AD biomarkers that carry unique predictive information rather than overlapping, redundant information. We operationalize this statistically by contrasting conditional with marginal importance. We exploit the accuracy of tabular foundation models [Hollmann et al., 2025; Qu et al., 2025] on multi-modal biomedical inputs to obtain well-powered importance estimates. To handle the pervasive correlations of high-dimensional neuroscience data, we present techniques for estimating group-level conditional importance. We focus in particular on proteomics data from blood plasma (PL) or cerebrospinal fluid (CSF) and neuroimaging data as these data modalities are being actively researched in the field of biomarker development in AD. We apply the CFI framework to three complementary large-scale cohorts: BioFINDER-1 (NCT01208675) [Palmqvist et al., 2021], a clinically specialized AD cohort with detailed CSF and plasma proteomics; UK Biobank [Sudlow et al., 2015], a large population-based study with broad plasma-proteomic coverage of thousands of analytes but no AD-specific focus; and ADNI [Jack Jr et al., 2008], which provides multimodal neuroimaging including magnetic resonance imaging (MRI) and positron emission tomography (PET). Across all cohorts, CFI identified protein and imaging signals providing unique and independent information on AD onset, diagnosis and progression whereas group-based CFI enabled statistical inference at the level of modalities, biological pathways and brain regions.

## 2 Results

### 2.1 CFI isolates predictive factors with unique information while controlling false discoveries

CFI is a versatile tool for measuring the unique predictive contribution of a feature within a multivariate model, in contrast to marginal (univariate) feature importance. CFI accommodates any ML model, is robust to correlations between variables and can be adapted to different prediction tasks, as shown in Figure 1 **a**. It targets a well-defined mathematical quantity: the total Sobol index, which generalizes the variance decomposition principle of ANOVA to non-linear models, quantifying the variance in the outcome attributable to a given variable [Homma and Saltelli, 1996; Williamson et al., 2021]. To achieve type-1 error control, CFI models the dependencies between variables by sampling the feature of interest from its conditional density (Figure 1 **a**.2). This generates a perturbed feature, which disrupts the unique information carried by that target feature while preserving the original correlation structure of the data. Consequently, measuring the performance drop between the initial input and the input containing the perturbed feature isolates the unique predictive contribution of the variable. This mechanism neutralizes the confounding effects of features that are merely correlated with important ones but do not add any unique information. In an extensive simulation benchmark comparing CFI with alternative methods, CFI effectively controlled type-1 error, even at high level of correlation, unlike explainability approaches such as traditional permutation feature importance (PFI) and SAGE (Figure 1 **b, c**). Furthermore, it maintained high statistical power (identifying truly important features) and outperformed model-X knockoff and DCRT which struggle with non-linear relationships, as shown in Figure 1 **d, e**, highlighting its capacity to uncover complex data-generating mechanisms. A presentation of the prior related work is provided in subsection 4.1, raw power values are reported in Figure S2. This combination of Type-1 error control, high statistical power, and interpretable magnitude estimation makes CFI suited for biomarker exploration in high-dimensional correlated data, as we demonstrate below.

**Figure 1:**
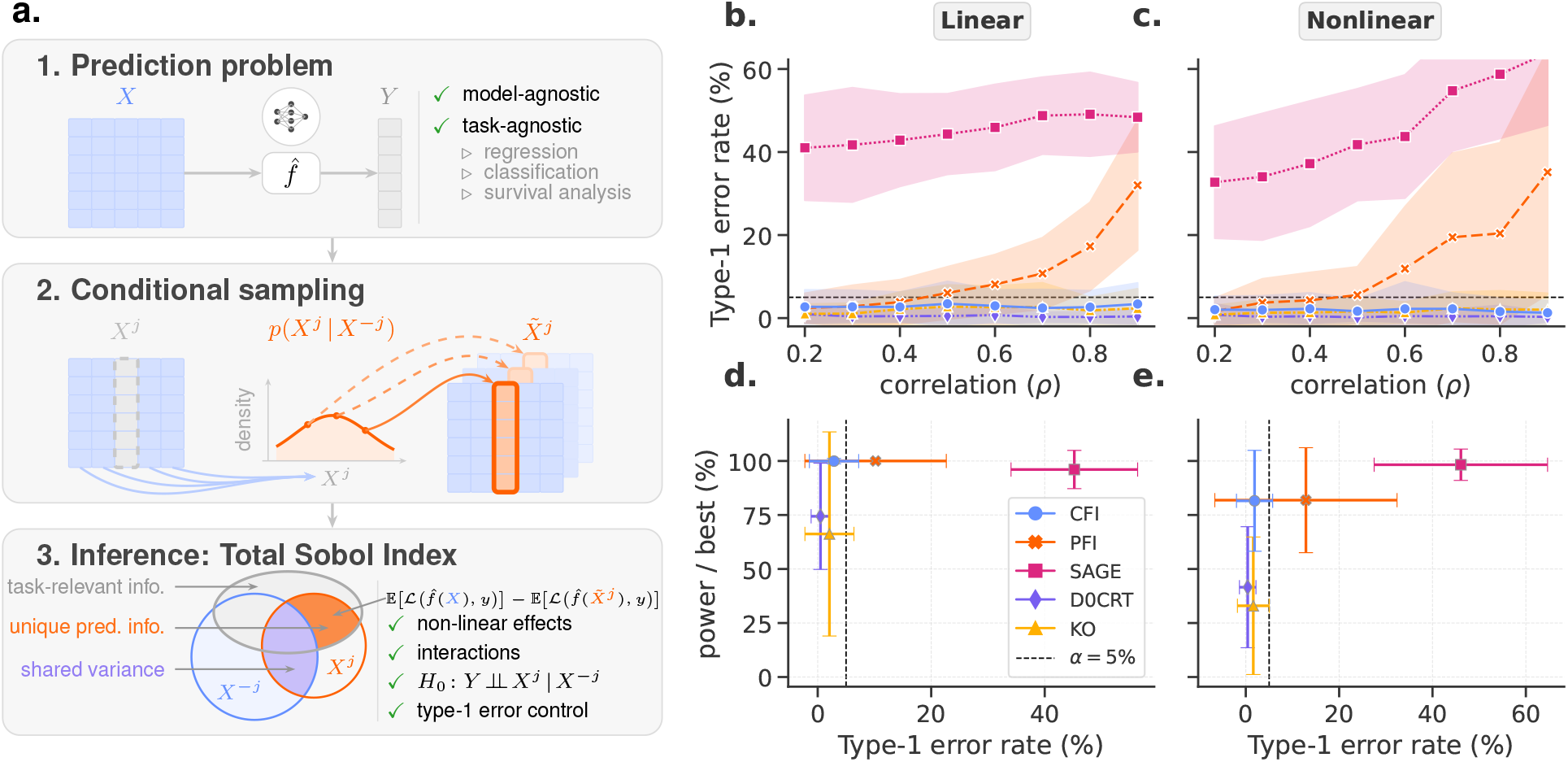
Conditional Feature Importance (CFI), a flexible tool for measuring importance with statistical guarantees. **(a)** The three stages of the CFI method. (1) *Prediction problem:* Fit a machine learning model *f* to predict a response *Y* from input features *X* for any standard task (e.g., regression, classification, survival analysis…). (2) *Conditional sampling:* Model the dependency between a feature of interest *X^j^* and the remaining features *X^−j^*, and generate a perturbed input *X̃* where the *j^th^* feature is sampled from the conditional distribution *p*(*X^j^ | X^−j^*). This step allows any arbitrary sampling method to be plugged in depending on the data type. (3) *Inference:* Compute the loss increase when predicting from the perturbed input *X̃* compared to the reference loss of *f* . This measure targets the total Sobol index (variance explained) and allows for conditional independence testing. Specifically, it tests against the null hypothesis that the feature provides no additional predictive value for the response when all other features are already known (*H*_0_ : *Y ⊥⊥ X^j^ | X^−j^*). **(b–e)** Benchmark comparisons of feature importance methods: Permutation Feature Importance (PFI), Shapley Additive Global importancE (SAGE), Distilled Conditional Randomization Test (DCRT), and Model-X Knockoffs (KO). Results are aggregated across 1000 simulated datasets with randomly sampled parameters, including the number of features, support sparsity (number of important features), sample size, feature correlation, and signal-to-noise ratio. **(b, c)** Type-1 error rates (nominal level *α* = 0.05, dashed line) as a function of feature correlation for a Ridge regressor in a linear scenario (**b**) and a neural network in a nonlinear scenario (**c**). **(d, e)** Statistical power divided by the power of the best method on each dataset (y-axis), versus Type-1 error rate (x-axis) for the linear (**d**) and nonlinear (**e**) scenarios. Error bars indicate one standard deviation.

### 2.2 Identification of non-redundant diagnostic and prognostic biomarkers in large AD cohorts

We first applied the CFI framework to identify which among strongly correlated protein biomarkers, genetic factors and demographic variables contain unique information for AD diagnosis and disease progression in BioFINDER. The analysis included 18 predictors: age, sex, *APOE ε*2 and *APOE ε*4 allele counts, 12 CSF biomarkers, and plasma neurofilament light-chain (NfL) and total tau. The CSF panel comprised A*β*_42_, A*β*_40_, total tau, p-tau181, NfL, GFAP, soluble TREM2 (sTREM2), YKL-40, *α*-synuclein, IL-6, neurogranin (NGN) and S100B. Between different machine-learning models, the more advanced tabular foundation models (TabICL and TabPFN) consistently outperformed established baselines like LightGBM (HGB) and regularized linear models (Figure 2 **a** and Figure S3). Therefore, we prioritized the tabular foundation models for the subsequent analyses. CFI then served to estimate the importance of the different features (proteomics in CSF and plasma [PL], demographics, genetics, the full list is provided in section 4.2.1) for both the diagnosis classification task (Figure 2 **b** left) and progression regression (Figure 2 **b** right). As CFI tests the null hypothesis that a given feature has zero conditional importance, this made it possible to infer whether a feature carries unique predictive information beyond all other features. Applied to AD protein biomarkers, it helped uncover differences between the variable importance profiles for AD diagnosis versus clinical progression. Baseline clinical diagnosis defined the three classes: 507 cognitively normal participants, 260 with mild cognitive impairment, and 47 with AD dementia. Among these variables, NGN, NfL, *α*-synuclein and A*β*_42_ in the CSF, total tau in plasma, age and biological sex rejected the null hypothesis of the conditional independence test after Benjamini-Hochberg (BH) multiplicity correction (*p <* 0.05), while CSF YKL-40 reached significance before correction. For the regression of the change in CDR-SB, age and CSF NfL, A*β*_42_, A*β*_40_ and p-tau181 remained significant after correction. Additional difference patterns emerged that were more subtle. CSF NGN and plasma total tau passed BH correction for diagnosis but not progression. CSF A*β*_40_ and p-tau181 passed BH correction for progression but not diagnosis. Age and CSF A*β*_42_ passed correction in both tasks.

**Figure 2:**
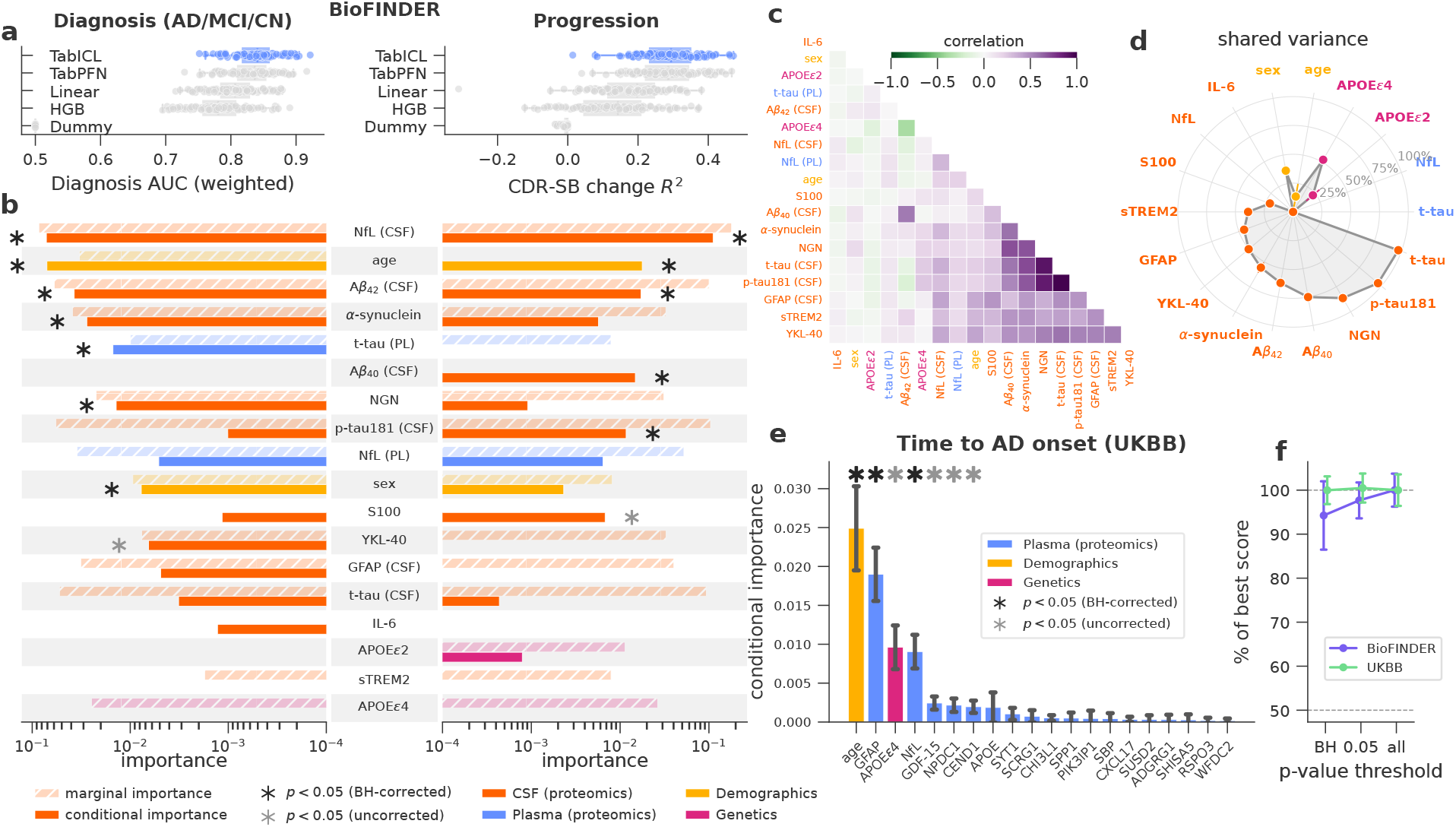
Conditional importance isolates unique predictive signals in correlated proteomic data. **(a)** Prediction benchmarks for diagnosis classification (ROC AUC) and CDR-SB regression (*R*^2^). The best-performing model (highlighted in blue) is used for measuring importance in (b). **(b)** Marginal and conditional importance of the top features for AD diagnosis (left) and CDR-SB progression (right) in BioFINDER. Faded hatched bars show the marginal importance, which quantifies the contribution of a feature irrespective of interactions and regardless of potential redundancies with other features. Solid bars show the conditional feature importance, which quantifies the unique predictive information of a feature, including possible interactions. Color encodes data modality. Asterisks mark significance of a conditional independence test (black: BH-corrected *p <* 0.05; gray: uncorrected). **(c)** Pearson correlation structure of the selected features. **(d)** Shared variance: percentage of each feature’s variance that can be explained by the remaining features. **(e)** Top proteins by conditional importance for a survival analysis of time to AD onset on UKBB proteomics data. **(f)** Percentage of best prediction score retained when selecting features at different p-value thresholds.

To contrast the unique predictive information obtained via CFI with uncorrected one-by-one association between the protein biomarkers and the AD diagnosis and progression, we also computed the marginal importance for each biomarker (Figure 2 **b**, hatched bars). The marginal importance was estimated using the Leave-One-Covariate-In (LOCI) procedure, which measures how much predictive performance increases above chance when a prediction model is trained on a single feature, providing a nonlinear but univariate measure of association. Because the marginal importance ignores interactions and does not account for redundancy among features, it can assign high importance to variables whose predictive signal is entirely shared with others. While most proteins carried predictive information, the comparison with CFI revealed that only a fraction carried unique information. A striking example is *APOE ε*4, the predominant genetic risk factor for late-onset AD. Despite a strong marginal importance, it carried no detectable conditional importance in BioFINDER, suggesting that the information it provides about AD risk was already contained in the measured protein biomarkers. This example of *APOE ε*4 is representative of a common observation in applied statistics: variables with a marginal importance, often identified in univariate analysis, do not necessarily show conditional effects, which is driven by redundancy between variables and vice-versa. This trend was visible across features in Figure S1, where the majority of variables lay above the identity line, indicating that their marginal association exceeded their unique conditional contribution.

However, the opposite pattern (conditional importance but no marginal importance) also appeared in our results. For instance, A*β*_40_ levels in the CSF showed no marginal importance but rather a strong conditional importance for progression. This suggests that CSF A*β*_40_ acts as a context variable: a feature with little predictive information on its own that helps interpret other biomarkers. This is consistent with the known biology of A*β*_40_, which reflects baseline inter-subject variability in amyloid production and CSF dynamics rather than pathology itself. A*β*_40_ is therefore routinely used to normalize A*β*_42_ levels, most commonly via the A*β*_42_/A*β*_40_ ratio, separating pathological A*β*_42_ reduction (due to plaque sequestration) from non-pathological variation. Age displays a similar profile, particularly for disease progression, where it has no marginal importance but strong conditional importance. This is consistent with age being a weak predictor of the rate of cognitive decline on its own, while being essential to interpret other biomarkers: the same level of tau or neurodegeneration carries a different prognosis depending on the patient’s age. Finally, while biological sex carried significant conditional importance for AD diagnosis, its conditional importance did not reach significance for disease progression. This asymmetry is consistent with the higher prevalence of dementia observed among women in aged populations [Knopman et al., 2021], although differences in recruitment across diagnostic groups may also contribute, and suggests that the contribution of sex to progression is weaker than its contribution to diagnosis.

To investigate the divergence between conditional and marginal importance, we computed the correlation matrix (Figure 2 **c**) and the shared variance (Figure 2 **d**) quantifying how well a feature can be predicted from all others. This analysis revealed that *APOE ε*4 genotype could be predicted from the remaining features with a cross-validated AUC of 0.761 (Figure 2d shows 2 *×* AUC *−* 1 = 0.522). It is noteworthy that the relationship between shared variance and conditional importance is not strictly negative and leaves room for subtlety. For example, a feature can have both a large shared variance and a high conditional importance if the remaining unexplained variance is highly predictive of the outcome. Despite sharing over 75% of its variance with other features, CSF A*β*_40_ remained highly conditionally important (BH-corrected p *<* 0.05), especially for predicting AD progression.

For the BioFINDER dataset, we also reported comparisons between CFI results across all machine learning models (Figure S4), showing high agreement between models. Moreover, we reported the commonly used permutation importance (Figure S5), showing a higher number of significant variables in line with its proven inability to control type-1 error in the presence of correlated data (see Chamma et al. [2023] and Figure 1 **b**-**e**).

To investigate scaling behavior of CFI with large datasets, we next turned our attention to AD-onset prediction within the UK Biobank (UKBB) cohort using nearly 3,000 plasma proteins across more than 46,000 participants with a regularized Cox proportional hazards model (Figure 2 **e**). Time-to onset labels were derived using the approach presented in Guo et al. [2024], using diagnoses from linked hospital, primary-care and death-registration records. The CFI analysis of the top 20 features highlighted similarities with diagnostic and progression prediction in BioFINDER, as well as some notable differences. In UKBB, age, plasma GFAP and plasma NfL rejected the conditional independence test after BH correction (*p <* 0.05). In BioFINDER, CSF NfL also passed correction, whereas CSF GFAP did not. However, *APOE ε*4 genotype was a key divergence, ranking third by conditional importance in UKBB (*p <* 0.05 before correction) after showing essentially none in BioFINDER. Moreover, additional proteins related to metabolic stress (GDF-15) and neuronal health (NPDC1, CEND1) reached significance before correction which have no counterpart in the BioFINDER panel. As these biomarkers track neurodegeneration but are not specific to AD pathology (in contrast to A*β* and phosphorylated tau proteins that were not part of the Olink panel), this may explain why the model relied on *APOE ε*4.

Comparing the large UKBB dataset to the smaller BioFINDER cohort further allowed to study the relationship between conditionally important features identified by CFI and model performance, testing whether CFI is capable of identifying a sufficient, reduced prediction model (Figure 2 **f** ). In UKBB, the curve was flat across thresholds: the features surviving BH correction were already sufficient to match the performance of the model using all features. In BioFINDER, only the uncorrected threshold reached full performance, while restricting to the BH-corrected set retained around 90% of it. With roughly 57 times more participants than BioFINDER (*n* = 46,382 versus *n* = 814), UKBB provides enough statistical power for the conditional independence test to recover every conditionally important feature even at the most stringent threshold, whereas in BioFINDER the BH correction discards features that are conditionally important but whose evidence is too weak at this sample size to survive multiplicity correction. Taken together, these results show that CFI identifies a reduced set of features that predicts as well as the full model: features with no unique conditional contribution can be discarded without loss, a property known as the sufficiency of the Markov blanket. As the contrast between the two cohorts illustrates, the stringency at which this reduced set can be selected inevitably depends on the sample size.

### 2.3 Group importance on highly correlated features in proteomics and neuroimaging

Previous work has highlighted that individual-level conditional importance mechanistically decreases if features are highly correlated [Verdinelli and Wasserman, 2024]. In the presence of high correlation, CFI leads to vanishing conditional importance which reduces the signal-to-noise ratio and thus statistical power. This principle also affects core AD biomarkers such as CSF total tau and p-tau181, which show substantial correlations. Moreover, genes are organized into pathways, giving rise to correlated sets of expressed proteins. In biomedicine and AD research, pathway-based methods therefore shift the focus from individual genes or proteins to biologically related groups. Likewise, in neuroimaging analyses, single voxels and even entire anatomical regions can be highly correlated and are thus grouped by researchers. This can be tackled with CFI by measuring the importance of variable groups instead of individual variables [Chamma et al., 2024].

The group-based CFI strategy retains the high correlation within each group to preserve statistical power while controlling type-1 error at the group level and can be extended to model the hierarchical relationship between variables [Paillard et al., 2025a]. Importantly, a key aspect of group feature importance is that the underlying predictive model does not reduce the variable set, which could degrade predictive performance, but instead uses every single feature inside each group, and thus swapping the null hypothesis *X^j^ ⊥ Y | X^−j^* for a feature *j*, to *X^G^ ⊥ Y | X^−G^* for a group *G*. Grouping is therefore applied when computing the feature importance, not the prediction model itself. Here, we applied the grouping approach to investigate importance of entire groups of proteomic and imaging features.

We first used data modality as the grouping criterion and analysed the conditional importance of plasma NfL and total tau, the same two markers in CSF, the remaining ten CSF biomarkers, the sociodemographic variables and the genotype for diagnosis classification in BioFINDER (Figure 3 **a**). The CSF panel excluding NfL and total tau carried the largest conditional group importance, followed by CSF NfL and total tau, demographics, and plasma NfL and total tau; all four groups were significant after BH correction. Comparing the same two markers across biofluids, NfL and total tau provided more conditional predictive information when measured in CSF than when measured in plasma. This is consistent with our biomarker-wise analysis (Figure 2 **b**), in which the most important individual markers came from CSF. The genetics block, consisting of *APOE ε*2 and *APOE ε*4 allele counts, did not turn out conditionally important.

**Figure 3:**
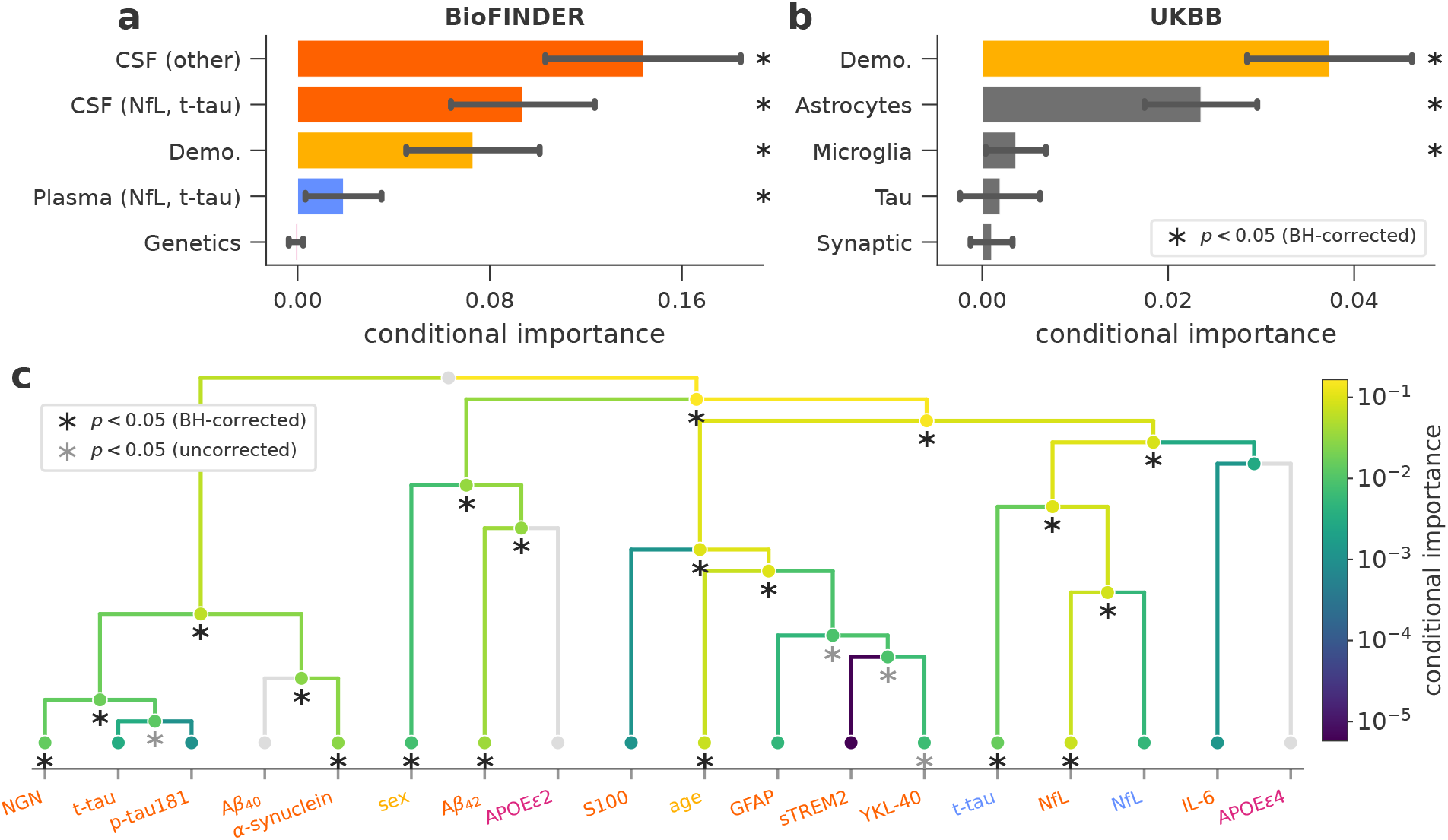
CFI identifies jointly predictive feature groups across data modalities, biological pathways, and correlation-based hierarchies. In all panels, grouping affects only the importance measurement: the predictive model always uses every feature. **(a)** Importance of feature groups for AD diagnosis classification in BioFINDER: the CSF panel excluding NfL and total tau, CSF NfL and total tau, plasma NfL and total tau, demographics, and genetics. Bars show the mean across ten repeats of 10-fold cross-validation and error bars the standard deviation. **(b)** Importance of feature groups defined by biological pathways for predicting time to AD onset in the UKBB, each group being the intersection between a gene set and the proteins measured by the Olink panel, with demographics as an additional group. **(c)** Multi-scale group importance along a Ward hierarchical clustering of the 18 BioFINDER features, for diagnosis classification. Each dot is a group of features tested as a whole, from single features at the leaves to all features at the root. Dot color encodes the mean conditional importance across ten repeats of 10-fold cross-validation on a logarithmic scale, each branch takes the color of the node below it, and light gray marks groups with null or negative estimated importance. Leaf labels are colored by modality as in **(a)**. In all panels, asterisks mark rejection of the conditional independence test (black: BH-corrected *p <* 0.05; gray: uncorrected), the correction being applied within each panel.

To study large numbers of correlated gene sets and proteins, pathway analysis is a popular modeling approach in computational biology. We used an enrichment analysis approach [Ekins et al., 2007] to study groups of biologically related proteins in the UKBB Olink proteomics panel for time-to-onset prediction (Figure 3 **b**), focusing in particular on groups of astrocytic, microglial, synaptic, and tau-related genes. Groups consisted of the intersection between gene sets and proteins measured in the UKBB Olink panel, listed in Table S1. The analysis revealed three significant blocks of variables: demographic, astrocytic proteins, and microglial proteins. The demographic group composed of age and sex was the most important for the model, followed by astrocytic proteins and, after a greater drop in importance, microglial proteins. Again, this analysis suggests that the significant pathways contained protein levels which are jointly predictive although they might share common variance.

To explore the breakdown of importance from top-level groups to single features, we applied the hierarchical group importance approach [Paillard et al., 2025a]. For simplicity, we focused on the BioFINDER dataset with its smaller protein panel and measured importance for diagnosis prediction (Figure 3 **c**), showing how CFI identifies groups of jointly predictive variables along a dendrogram. The analysis revealed that conditional importance increased monotonically with group size, the root of the hierarchy carrying the largest importance and corresponding to the total variance explained by the model, while importance progressively decreased as groups were split into smaller and more correlated subsets. Then, descending along the tree, we observed that the importance of some groups could not be broken down to the level of each unique variable due to their high correlation. For instance, established AD biomarkers such as CSF total tau and ptau181 clustered together, alongside NGN, consistent with their biological proximity. This group carried significant conditional importance, although total tau and p-tau181 were not individually significant. The analysis also revealed clusters of variables related to distinct biological processes, including a cluster containing age and CSF GFAP, sTREM2 and YKL-40. Moreover, we observed that measuring the same protein across different biofluids (CSF and plasma) was not necessarily redundant, in line with the complementary role of CSF vs plasma (Figure 3 **a**) seen above. Specifically, CSF and plasma levels of total tau merged only near the top of the dendrogram, reflecting low inter-fluid correlation. This divergence may stem from peripheral protein sources or complex transfer dynamics between biofluids.

We next applied grouped importance to study AD diagnosis prediction for multimodal brainimaging data from the ADNI dataset (Figure 4). Here, grouping is not only useful for assessing the importance of different imaging modality i.e. MRI and PET but also probing the importance of spatially correlated signals across the brain within each modality. We predicted AD diagnosis (CN, MCI, AD) using TabICLv2 [Qu et al., 2026], a tabular foundation model, using multimodal neuroimaging data for in-context learning. The features included cortical thickness in 68 regions of the Desikan–Killiany atlas, volume in subcortical areas, amyloid levels in the cortical regions and five bilateral subcortical structures, age and biological sex. In a first step, our CFI analysis uncovered distinct patterns of unique predictive information conveyed by MRI cortical thickness and amyloid PET with both imaging modalities showing large conditional importance and significance (p-value *<* 0.05 after BH correction, Figure 4 **a**). Demographic features were also significant, but with much lower conditional importance.

**Figure 4:**
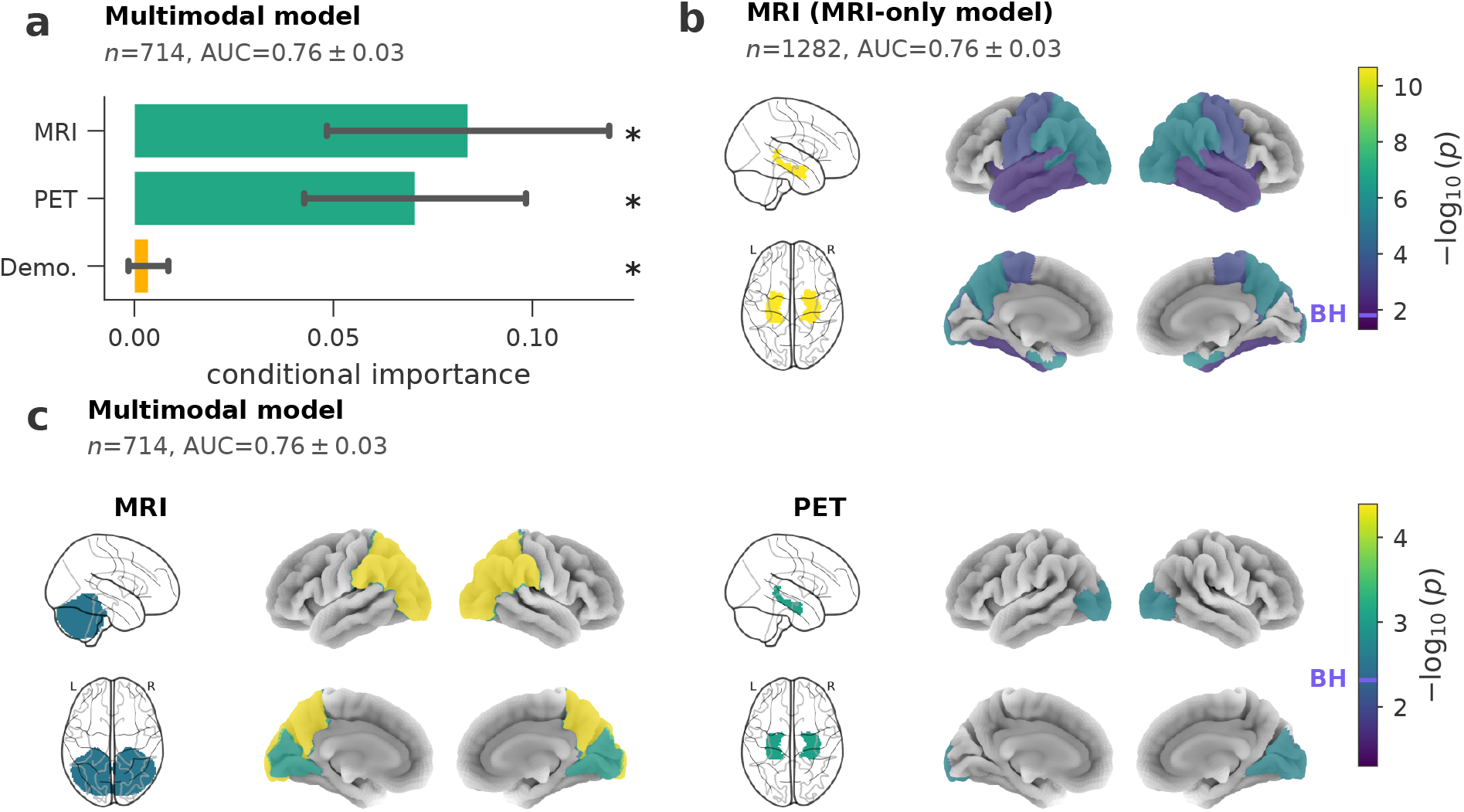
CFI reveals complementary signals across neuroimaging modalities and highlights brain regions carrying unique predictive information. Group-level conditional importance for AD diagnosis (CN, MCI, AD) in ADNI, using a TabICL classifier and 10-fold cross-validation. Both models predict from individual regions: cortical thickness in the Desikan– Killiany (DK) atlas, subcortical volumes and demographics for the MRI-only model, to which the multimodal model adds amyloid PET SUVRs in the DK regions and subcortical structures; grouping enters only when importance is measured, and no model is refitted. Sample size and cross-validated ROC AUC are indicated in each panel; the multimodal model is restricted to the participants with both scans. **(a)** Importance of the group containing all features of a modality, in the multimodal model, with bars showing conditional importance, error bars one standard deviation over the 10 folds, and asterisks significance after BH correction at *α* = 0.05. **(b)** Brain map of the *p*-values obtained for the MRI-only model. Importance is computed for 15 clusters of regions obtained by unsupervised clustering (clusters presented in Figure 5), every region taking the value of its cluster. **(c)** Same as **(b)** for the multimodal model, mapped for MRI (left) and PET (right); adding PET importantly changes the conditioning set, MRI importance now being conditional on PET as well. Color encodes *−* log_10_(*p*) and clusters below an uncorrected *α* = 0.05 are left uncolored; **(b)** has its own color scale, while the two maps of **(c)** share one. In **(b)** and **(c)**, subcortical structures are shown as glass-brain projections (left, sagittal above axial) and the cortical surface projected on *fsaverage* (right, lateral above medial, left hemisphere on the left). Purple marks on the colorbars indicate the BH significance thresholds.

To explore regional importance, we used group-based CFI with data-driven clustering, separately for PET and MRI and measured importance at the group level. Within each modality, we grouped regions of interest (ROIs) into 15 clusters using Ward’s clustering (Figure 5) and computed CFI at the cluster level for AD diagnosis in ADNI (Figure 4 **b** and **c**). Importantly, clusters were only applied at the importance step: the model was built using the individual ROIs from MRI and PET without dimensionality reduction at any point. We first measured importance using only MRI and demographics, from a cohort of *n* = 1282 participants (Figure 4 **b**), the AUC for the three-way classification problem was 0.76. Conditional importance recovered the established structural hallmarks of AD: bilateral hippocampal and amygdalar volume carried by far the strongest unique signal (*p* = 2.2 *×* 10*^−^*^11^), followed by a bilateral posterior-parietal cortical-thickness cluster spanning the precuneus, superior and inferior parietal, supramarginal, lateral occipital cortex and the banks of the superior temporal sulcus (*p* = 1.3*×* 10*^−^*^5^), both significant after BH correction over the 16 clusters tested. Entorhinal and temporal-pole thickness (*p* = 7.5 *×* 10*^−^*^6^) and a cluster spanning the paracentral, precentral, postcentral and transverse temporal cortices (*p* = 0.0038) were also significant after correction. We then measured importance in a multi-modal model, predicting from MRI, amyloid PET and demographics which achieved an AUC=0.76 on the cohort of *n* = 714 participants with both scans. Adding amyloid PET redistributed this signal rather than adding to it (Figure 4 **c**): once amyloid entered the conditioning set, hippocampal volume no longer carried any unique information. Instead, the hippocampal signal was captured by amyloid levels in the same structure (*p* = 0.029), the only PET cluster with significant conditional importance. Because the model included both PET and MRI-derived regional volumes, hippocampal PET importance was assessed conditional on these volumes, controlling for measured anatomical differences relevant to partial-volume effects. Posterior-parietal thickness (*p* = 4.1 *×* 10*^−^*^5^) and occipital thickness in the cuneus, lingual and pericalcarine cortices (*p* = 0.0037) remained significant, with both MRI clusters and the hippocampal PET cluster surviving BH correction across the 31 clusters of this multimodal analysis. These results demonstrate the capacity of CFI for identifying biologically relevant patterns from highly correlated neuroimaging features that individual-level analysis cannot resolve.

**Figure 5:**
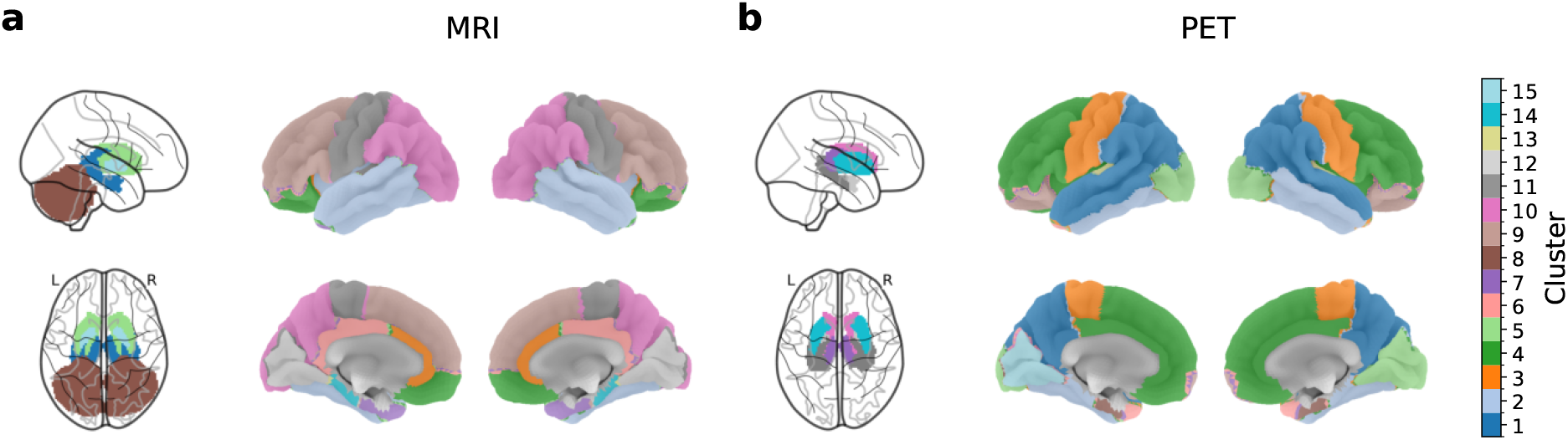
Ward clustering of neuroimaging ROIs. **(a)** MRI cortical thickness and **(b)** A*β* PET regions of interest from the Desikan–Killiany atlas (cortical) and AAL atlas (subcortical), colored by cluster membership. Ward’s agglomerative clustering with 15 clusters per modality was applied to the feature matrix, grouping ROIs with similar profiles across subjects. These clusters define the groups used to compute the conditional importance maps shown in Figure 4 **b** and **c**. Left panels show subcortical structures (sagittal and axial glass brain projections); right panels show cortical surfaces (lateral and medial views).

## 3 Discussion

In this work, we presented CFI, a framework for statistical inference with any predictive ML model based on estimating conditional variable importance. We applied CFI to the problem of identifying unique predictive signals of AD inside large proteomics and imaging datasets characterized by substantial overlap and redundancy between biomarker candidates. We demonstrated the ability of CFI to leverage state-of-the-art tabular foundation models and its versatility across different prediction tasks (diagnosis, progression, time to onset) on three large AD cohorts (BioFINDER, ADNI, UKBB) encompassing diverse data modalities: proteomics, demographics, MRI, and A*β*-PET. Importantly, CFI allowed us to identify the unique predictive information of biomarkers and sociodemographic factors by contrasting conditional with marginal importance. This was illustrated by *APOE ε*4, which has a strong marginal association but no conditional importance when key proteins such as amyloid and tau were accounted for. Furthermore, it highlighted the independent information captured by NfL, *α*-synuclein and NGN in CSF against the background of multiple redundant biomarkers, as well as groups of proteins defined in terms of gene pathways. Moreover, CFI consistently scored A*β*-PET and MRI as complementary for AD diagnosis, providing insights into regional cortical and subcortical localization of information. Our results highlight the practical value of the CFI framework for parsing statistically independent prediction signals from large multimodal datasets in biomedicine.

### The CFI framework provides inference on correlated biomarker data in AD

CFI targets the Total Sobol Index [Homma and Saltelli, 1996], which is the best quantity for identifying the smallest set of predictive features in the presence of correlation [Bènard et al., 2022] and thus enables inference with statistical guarantees. This theoretical property was empirically validated, in a benchmark showing that CFI controls the risk of false discoveries even at high levels of correlation, contrary to alternative feature importance methods (Figure 1). Applied to AD diagnosis and prognostic prediction across three large-scale cohorts, CFI revealed distinct variable roles that align with known biology. Age and NfL were important for all three outcomes, and A*β*_42_ was common to diagnosis and progression. Beyond this core, in BioFINDER, diagnosis was specifically predicted by total tau (plasma), *α*-synuclein, neurogranin (NGN), sex, and, uncorrected, YKL-40, whereas progression was specifically predicted by p-tau181, A*β*_40_, and, uncorrected, S100. In UKBB, plasma GFAP was specifically linked to onset. These different signatures potentially reflect cumulative neuronal and synaptic injury, alongside sex differences, in diagnosis versus active amyloid-cascade signaling for progression, whereas GFAP may capture inflammatory signaling reflecting early disease activity.

More broadly, these results provide key distinctions between features with high marginal association but no unique conditional contribution, and conversely, features with no marginal association but strong conditional importance. The most prominent example of the former is *APOE ε*4, the predominant genetic risk factor for late-onset AD, which carries near-zero conditional importance despite a strong marginal importance. This suggests that its predictive contribution in BioFINDER overlaps with information captured by the measured proteins, consistent with *APOE ε*4’s role as an upstream modulator of amyloid and tau pathology and its associated proteomic changes [Lu et al., 2026]. Supporting this interpretation, *APOE ε*4 regains high conditional importance in the UKBB cohort, where the Olink panel does not include AD-specific biomarkers such as A*β* or phosphorylated tau proteins and, therefore, cannot explain away its signal. Conversely, CSF A*β*_40_ exemplifies the opposite pattern: it exhibits a negligible marginal importance but strong conditional importance, consistent with its known role as a reference for baseline amyloid production: A*β*_40_ levels are uninformative on their own but necessary to determine whether a given A*β*_42_ level reflects pathological accumulation or normal inter-subject variability. CSF total *α*-synuclein also carried predictive information for diagnosis conditional on the other measured variables. Its correlations with tau, NGN and A*β*_40_ (Figure 2c) suggest a contribution related to neuronal injury and shared variation in CSF protein concentrations. Its usefulness within a multivariable panel is consistent with the improved differential diagnosis reported when combining total *α*-synuclein with amyloid and tau biomarkers by Shi et al. [2018]. Importantly, CFI also helped distinguish features whose roles differ across clinical endpoints. NGN, for instance, carried significant conditional importance for AD diagnosis but not for disease progression. Beyond the classical ATN (Amyloid, Tau, Neurodegeneration) axes, CFI identified significant conditional predictive information in plasma GFAP in UKBB, with additional evidence for CSF YKL-40 in BioFINDER before multiplicity correction. Conversely, other related markers like sTREM2 only showed marginal importance, which is in line with the capacity of CFI to isolate the strongest unique signal among multiple overlapping signals. Closely related, the astrocyte pathway further emerged as the most important genetically defined biological pathway group of proteins in the UKBB analysis. These findings are consistent with a growing body of work suggesting that neuroinflammation constitutes a fundamental axis of AD pathology beyond the classical amyloid, tau, and neurodegeneration (ATN) framework [Long and Holtzman, 2019]. The metabolic (GDF15) and neurosynaptic proteins (CEND1 and NPDC1) found in analysis of the UKBB plasma proteomics data were statistically less prominent but argue in the same direction beyond the ATN framework.

### CFI enables statistical inference with tabular foundation models

Tabular foundation models (TFMs) [Hollmann et al., 2025; Qu et al., 2025, 2026] have recently emerged as leading predictors for tabular data, a domain previously dominated by tree-based models [Grinsztajn et al., 2022]. Here, we demonstrated their ability to predict AD diagnosis and progression from complex, multimodal, high-dimensional data, and showed that CFI can harness the predictive power unlocked by their large-scale pretraining. By identifying the features on which these models base their predictions, CFI makes their otherwise opaque predictions interpretable and, unlike alternative attribution methods, provides statistical guarantees for the resulting importance scores. These insights can guide the identification of candidate diagnostic biomarkers or generate hypotheses about potential therapeutic targets. Although such hypotheses can be generated computationally, validating them requires costly collection of new data. The valid p-values provided by CFI help researchers to control the risk of pursuing misleading leads and prioritize candidates for further investigation.

### Implications for biomarker development

The combination of strong prediction models with statistically sound inference rules has several implications for biomarker discovery and development. Discoveries made with CFI prevent redundancy, as the set of conditionally important features forms a sufficient set for optimal prediction (Figure 2 (**f** )), avoiding the inclusion of variables whose signal is already captured by others. This can help generate hypotheses that, for example, inform the development of surrogate biomarkers in the context of clinical trials, e.g. when scanning through a large protein panel to detect biomarker changes that predict clinical efficacy. Such biomarkers can provide evidence of a therapeutic effect before the readout of clinical endpoints, which typically involves much longer time scales. This is for instance the case in AD, where changes in cognitive scores take years to manifest while biomarkers such as amyloid and tau provide earlier evidence of a potential therapeutic effect. Given the large number of biomarkers implicated in AD pathophysiology, selecting a parsimonious set is essential to limit patient burden and clinical trial costs. CFI provides a principled approach to this problem by framing biomarker selection as conditional importance testing, addressing key limitations of existing approaches. For instance, Oh et al. [2025] used recursive feature elimination to identify the CSF YWHAG and NPTX2 proteins and quantified their importance for predicting cognitive impairment using coefficients in a Lasso model. However, this approach is specific to linear models and requires hand-crafting non-linear transformations such as protein ratios to capture non-additive effects, whereas CFI can leverage more flexible models that learn such relationships directly from the data. In addition, CFI provides statistical guarantees, which are crucial for decision making that recursive methods cannot achieve. In large-scale proteomic studies of the UKBB, using a similar recursive approach, Guo et al. [2024] and Deng et al. [2025] both identified GFAP, NfL, and GDF15 among the top proteins associated with dementia prediction, consistent with the results of CFI. However, because neither approach controls the risk of false discoveries, both studies also retain many additional proteins that may carry redundant information rather than unique predictive contributions. In practice, this leads to the inclusion of redundant biomarkers, as evidenced by a performance plateau in their forward selection curve: predictive accuracy saturates after a handful of proteins while additional ones continue to be selected. Such redundancy is a direct concern for clinical translation, where each candidate biomarker must be validated experimentally, and pursuing markers that provide no information beyond what is already captured by others wastes resources and increases patient burden. As an alternative, Mi et al. [2021] presented a method for the identification of biomarkers based on PFI. PFI importances on BioFINDER are shown in Figure S5. While allowing to leverage any ML model, it lacks statistical guarantees and inevitably leads to false discoveries at high levels of correlation (as shown in Figure 1, see also Chamma et al., 2023). CFI addresses this by providing p-values that control false discoveries, ensuring that only biomarkers carrying unique predictive information are retained.

Furthermore, the machine-learning pipelines commonly applied to large datasets, e.g. Guo et al. [2024] and Deng et al. [2025], perform feature selection (often via univariate Cox proportional hazard models for AD risk and volcano plots), model fitting, and inference as separate sequential steps, a design prone to data leakage: when selection uses the outcome outside the cross-validation loop, both the reported performance and the resulting p-values are over-optimistic. By contrast, our proposed pipeline (c.f. Figure 2 **e**) unifies these operations into a single statistically controlled pipeline by running Cox proportional-hazard machine-learning model with built-in feature selection procedure and CFI for variable importance analysis.

### Extension to causal inference and treatment effect heterogeneity

Beyond predicting disease outcomes, predicting how treatment effects vary across patients is a central problem in drug development. Identifying which patient characteristics modulate treatment effects can improve the probability of success of clinical trials by targeting the right population, accelerate timelines by pinpointing likely responders, and reduce costs by avoiding enrollment of patients unlikely to benefit [Ruhrberg Estèvez et al., 2026]. Estimating conditional average treatment effects (CATE) from biomarker data is an essential step in the discovery of *predictive biomarkers*. This is particularly relevant in AD, where treatment effects can be highly heterogeneous: in the EMERGE trial, the benefits of *aducanumab* in terms of slowing cognitive decline varied greatly within the treated population [Pang et al., 2023]. Understanding what drives such variability could help identify subgroups of better responders, or define exclusion criteria based on risk factors for adverse events. CATE estimation goes beyond a single average treatment effect to characterize how treatment response varies across patient profiles (at the cost of assumptions, such as the absence of confounders, the overlap of populations in treatment arms, that are hard to verify in practice). CFI can be extended to this setting by measuring feature importance in CATE models, identifying treatment-effect modifiers while retaining the same guarantees for false-discovery control [Paillard et al., 2025b; Abècassis et al., 2025].

### Broader applicability

The present work focuses on AD, motivated by the intrinsic complexity of the disease: AD is multifactorial, polygenic, encompasses heterogeneous clinical subtypes, and involves interacting biological processes. These properties make AD a particularly demanding testbed, yet they are by no means unique to this disease. Indeed, CFI has been applied to other biomedical problems, including the identification of genes characterizing breast cancer subtypes, proteomic signatures of body mass index and lifestyle habits, and changes in cortical activity measured with EEG and MEG [Paillard et al., 2026; Chamma et al., 2023; Gaubert et al., 2025; Paillard et al., 2025a]. More broadly, CFI extends naturally to any supervised learning problem, including domains outside biomedicine.

### Limitations

A limitation of CFI is that the insights it generates are only as good as the underlying model. Indeed, if the model fails to adequately capture the response, the resulting importance decomposition will be unreliable. However, tabular foundation models, which leverage large-scale pretraining, have been shown to achieve strong predictive performance even on moderately sized datasets, partially mitigating this concern. Another limitation is computational: measuring feature importance requires fitting at least one additional model [Hooker et al., 2021]. Because CFI trains one conditional sampler per feature, this cost scales with dimensionality and can become prohibitive in high-dimensional settings. Grouping addresses this directly, one sampler per group rather than per feature, and simultaneously restores the power lost to strong correlation, but it displaces the difficulty onto the choice of grouping. A pre-specified grouping requires domain knowledge that is costly or simply unavailable. On the other hand, data-driven grouping requires specifying the number of clusters, therefore introducing a trade-off between the resolution of the inference and the statistical power. While larger groups improve statistical power, they provide a coarser inference which is arguably less informative. The hierarchical approach attempts to solve this challenge.

## Data availability

Individual-level data are subject to controlled access. BioFINDER-1 data may be requested from the Swedish BioFINDER study group, subject to applicable ethical approvals and a data-transfer agreement. ADNI data are accessible through the LONI Image and Data Archive following access approval. UK Biobank data access is governed by its application and approval procedures. This study used UK Biobank application 44257.

## Code availability

The implementation of CFI along with other baseline methods studied in this paper is available in hidimstat^1^, an open-source library with full compatibility with the scikit-learn API which makes it modular and flexible allowing the use of any ML model that is also compatible with the scikitlearn API. The experimental code specific to this manuscript will be made publicly available upon publication.

## Acknowledgments

This research has been conducted using the UK Biobank Resource under Application Number 44257. We thank the participants and investigators of the Swedish BioFINDER study for providing the data used in this work. A complete list of the BioFINDER study group members is available at https://www.biofinder.se. BioFINDER research at Lund University received support from the Swedish Research Council (2022-00775), ERA PerMed (ERAPERMED2021-184), Knut and Alice Wallenberg Foundation (2017-0383), MultiPark at Lund University, Swedish Alzheimer Foundation (AF-980907), Swedish Brain Foundation (FO2021-0293), Parkinson Foundation of Sweden (1412/22), Cure Alzheimer’s Fund, Konung Gustaf V:s och Drottning Victorias Frimurarestiftelse, Skåne University Hospital Foundation (2020-O000028), Regionalt Forskningsstöd (2022-1259), and the Swedish government through the ALF agreement (2022-Projekt0080).

Data collection and sharing for the Alzheimer’s Disease Neuroimaging Initiative (ADNI) is funded by the National Institute on Aging (National Institutes of Health Grant U19 AG024904). The grantee organization is the Northern California Institute for Research and Education. In the past, ADNI has also received funding from the National Institute of Biomedical Imaging and Bioengineering, the Canadian Institutes of Health Research, and private sector contributions through the Foundation for the National Institutes of Health (FNIH) including generous contributions from the following: AbbVie, Alzheimer’s Association; Alzheimer’s Drug Discovery Foundation; Araclon Biotech; BioClinica, Inc.; Biogen; Bristol-Myers Squibb Company; CereSpir, Inc.; Cogstate; Eisai Inc.; Elan Pharmaceuticals, Inc.; Eli Lilly and Company; EuroImmun; F. Hoffmann-La Roche Ltd and its affiliated company Genentech, Inc.; Fujirebio; GE Healthcare; IXICO Ltd.; Janssen Alzheimer Immunotherapy Research & Development, LLC.; Johnson & Johnson Pharmaceutical Research &Development LLC.; Lumosity; Lundbeck; Merck & Co., Inc.; Meso Scale Diagnostics, LLC.; NeuroRx Research; Neurotrack Technologies; Novartis Pharmaceuticals Corporation; Pfizer Inc.; Piramal Imaging; Servier; Takeda Pharmaceutical Company; and Transition Therapeutics.

## Author contributions

Conceptualisation : JP, JB, TKT, ARL, AC, BT, DE. Methodology: JP, ES, JB, TKT, ARL, BT, DE. Software and formal analysis: JP, ES, AC, BT. Data curation: JP, ES, JB, TKT, NMC. Visualization: JP, BT, DE. Writing – original draft: JP, BT, DE. Writing – review & editing: JP, ES, JB, TKT, GDP, ARL, AC, ES, SP, SJ, NMC, BT, DE. Supervision: BT, DE.

## Competing interests

JP, ES, JB, TKT, GDP and DE, are employees of F. Hoffmann-La Roche Ltd. NMC has received speaker/consultancy fees from BioArctic, Biogen, BMS, Eli Lilly, Genentech, Merck, Novo Nordisk, Owkin, Positrigo and Roche. SP has acquired research support (for the institution) from Avid Radiopharmaceuticals and ki elements through ADDF. He has received consultancy/speaker fees from BioArctic, Biogen, Danaher, Eisai, Eli Lilly, Novo Nordisk, Roche, and Sanofi. ES has acquired research support (for the institution) from Beckman Coulter, Bristol Myers Squibb, C2N Diagnostics, Eisai, Fujirebio, GE Healthcare and Roche Diagnostics.

## 4 Methods

### 4.1 Related work

Several model-agnostic approaches have been proposed to tackle estimation of variable importance inside prediction models each of which faces specific limitations. Existing model-agnostic variable importance measures like Permutation Feature Importance (PFI) [Breiman, 2001] and Shapley Additive Global importancE (SAGE) [Covert et al., 2020] quantify feature importance across any machine learning model. However, they lack statistical guarantees, inevitably yielding false discoveries in the presence of correlated features (Figure 1). Conversely, conditional independence tests such as model-X knockoffs [Candes et al., 2018] and distilled conditional randomization testing (DCRT) [Liu et al., 2022] perform statistically rigorous variable selection while all other variables are already accounted for. However, for knockoffs, the most powerful approach relies on the Lasso Coefficient Difference (LCD) statistic which can only identify linear effects, and DCRT is limited to additive effects, missing any predictive signal that arises through interactions between variables. Moreover, both are variable selection methods rather than variable importance measures; neither quantifies the magnitude of each variable’s contribution. Bridging this gap requires measuring conditional feature importance to simultaneously quantify a feature’s impact and provide formal statistical guarantees. The primary method achieving this is ablation, also known as Leave-One-Covariate-Out (LOCO) [Williamson et al., 2023]. However, LOCO requires computationally prohibitive model refitting for every feature, which also makes the estimation noisy. Conditional Feature Importance (CFI) approaches target the same mathematical quantity as LOCO but bypass its computational burden, while also outperforming it in statistical power [Chamma et al., 2023; Paillard et al., 2025b]. We describe below the datasets, the CFI framework and its implementation, the statistical testing procedure, and the feature grouping strategies used throughout this study.

### 4.2 Datasets

We applied CFI to three complementary cohorts spanning AD-specific proteomics (BioFINDER-1), large-scale population proteomics (UK Biobank), and multimodal neuroimaging (ADNI), selected to test the method across diverse data modalities, scales, and prediction tasks.

#### 4.2.1 BioFINDER-1

We used data from the BioFINDER-1 study (NCT01208675) [Palmqvist et al., 2021], comprising *n* = 814 participants with baseline cerebrospinal fluid (CSF), plasma, genetic, and demographic measurements. The analysis included 18 predictors: 12 CSF markers, plasma NfL and total tau, age, sex, and *APOE ε*2 and *APOE ε*4 allele counts. CSF biomarkers were measured using Roche Elecsys NeuroToolKit immunoassays Johnson et al. [2023]. Plasma analyses were restricted to NfL and total tau, with the same predictor panel used for diagnosis and progression. The same predictor panel was used for diagnosis and progression. For the diagnostic classification task, participants were categorized into three groups: cognitively normal (CN), mild cognitive impairment (MCI), and Alzheimer disease (AD). For the disease progression task, we modeled longitudinal change in the Clinical Dementia Rating Scale Sum of Boxes (CDR-SB), a primary or key secondary endpoint in recent clinical trials of amyloid-lowering therapies [Van Dyck et al., 2023; Sims et al., 2023], using a linear mixed-effects model with fixed effects for baseline age, sex, baseline CDR-SB, a diagnosisby-timepoint interaction, and a random slope per participant (intercept constrained to zero). The progression analysis included 724 participants, of whom 484 were cognitively normal and 240 had MCI at baseline. Covariate-adjusted individual progression rates were extracted as participantspecific random slopes and rank-normalized. Participants with fewer than two longitudinal visits were excluded.

#### 4.2.2 UK Biobank

We used proteomic data from the UK Biobank [Sudlow et al., 2015], comprising *n* = 46,382 participants with measurements from the Olink Explore panel, yielding 2,922 plasma protein levels after quality control. Olink normalized protein expression (NPX) data were analyzed without further preprocessing, following the processing described in the UK Biobank Pharma Proteomics Project documentation (filed id 4658). Additional features included *APOE ε*4 allele count, age, and sex. The prediction target was time from protein measurement to AD diagnosis, modeled as a survival analysis endpoint. Participants without an AD diagnosis were right-censored at their last available follow-up.

#### 4.2.3 ADNI

Data used in the preparation of this article were obtained from the Alzheimer’s Disease Neuroimaging Initiative (ADNI) database (adni.loni.usc.edu). The ADNI was launched in 2003 as a public-private partnership, led by Principal Investigator Michael W. Weiner, MD. The primary goal of ADNI has been to test whether serial magnetic resonance imaging (MRI), positron emission tomography (PET), other biological markers, and clinical and neuropsychological assessment can be combined to measure the progression of mild cognitive impairment (MCI) and early Alzheimer’s disease (AD).

We used data from the Alzheimer’s Disease Neuroimaging Initiative (ADNI), comprising *n* = 714 participants with both structural MRI and ^18^F-florbetapir (AV-45) amyloid PET imaging. The feature set consisted of 164 variables: 2 demographic features (age, sex), 84 structural MRI features, and 78 amyloid PET features. MRI features comprised cortical thickness in the 68 regions of the Desikan–Killiany (DK) atlas [Desikan et al., 2006] (34 per hemisphere) and the volumes of 8 bilateral subcortical structures (amygdala, caudate, cerebellum cortex, cerebellum white matter, hippocampus, pallidum, putamen, and thalamus). Amyloid PET features comprised regional SUVRs from the 68 cortical DK regions and 5 bilateral subcortical structures (amygdala, caudate, putamen, hippocampus, and thalamus). Pallidum was excluded due to known quantification artifacts, and cerebellar regions were excluded, as the whole cerebellum served as the reference region for SUVR normalization. T1-weighted structural MRI was processed using FreeSurfer 7.3.2 recon-all [Fischl, 2012] for cortical reconstruction and parcellation. Cortical thickness was extracted for each region of the DK atlas. Pre-processed ^18^F-florbetapir (AV-45) PET images (co-registered, averaged, and smoothed to a uniform resolution) were downloaded from the ADNI LONI database. Each image was registered to the participant’s FreeSurfer-processed T1-weighted scan using mri coreg, and regional standardized uptake value ratios (SUVRs) were computed by normalizing mean up-take in each Desikan–Killiany region to the whole cerebellum. Early and late MCI were combined into a single MCI group for the three-way diagnostic classification (CN vs. MCI vs. AD).

### 4.3 Conditional Feature Importance

#### 4.3.1 Estimand

Let *f* denote a predictive model trained to predict a response *Y* from a feature vector *X* = (*X*^1^*, . . . , X^p^*), and let *ℓ* be a loss function. We define the perturbed input *X̃*(*^j^*) as the vector obtained by replacing the *j*-th component of *X* with a sample from its conditional distribution: *X̃* (*^j^*) = (*X*^1^*, . . . , X^j−^*^1^*, X̃^j^, X^j^*^+1^*, . . . , X^p^*) where *X̃^j^ ∼ p*(*X^j^ | X^−j^*). This construction preserves the dependencies between feature *j* and the remaining features while breaking its relationship with the output *Y* . The conditional feature importance of *X^j^* is defined as the expected loss increase nder this perturbation:

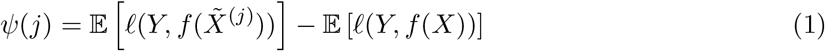

This quantity is proportional (up to a factor 2) to the total Sobol index of *X^j^* [Homma and Saltelli, 1996; Williamson et al., 2021] and it equals zero if and only if *X* is not predictive of the outcome *Y* given the rest. For instance, if *Y ⊥⊥ X^j^ | X^−j^*, therefore enabling a formal conditional independence test. For classification tasks, we used the cross-entropy loss function. For regression tasks, we used mean squared error *ℓ*(*y, f* (*x*)) = (*y − f* (*x*))^2^. For survival analysis (UK Biobank), we used the concordance index.

#### 4.3.2 Statistical testing and Inference

CFI yields, for each feature *j* and each cross-validation fold *k*, an importance estimate *ψ*^^^*_k_*(*j*). Under the null hypothesis *H*_0_ : *Y ⊥⊥ X^j^ | X^−j^*, the population importance *ψ*(*j*) = 0, and the per-fold estimates fluctuate around zero. We tested *H*_0_ using a one-sided *t*-test with the Nadeau–Bengio variance correction [Nadeau and Bengio, 1999], which accounts for the dependence between crossvalidation folds arising from overlapping training sets. Specifically, because cross-validation folds share training data, the per-fold estimates are positively correlated, making the naive variance of the mean (*σ*^^2^*/K*) overly optimistic. The correction inflates this estimate to 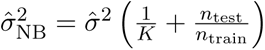 where *K* is the total number of held-out fold evaluations across repeats, *σ*^^2^ the sample variance across folds, and *n*_test_*/n*_train_ the test-to-train ratio [Nadeau and Bengio, 1999]. The resulting *t*-statistic follows a *t*-distribution with *K −* 1 degrees of freedom under *H*_0_. We used a one-sided test (alternative: *ψ*(*j*) *>* 0), as negative importance values are not meaningful under the loss-difference formulation. To control the false discovery rate across the *p* features tested simultaneously, *p*-values were corrected using the Benjamini–Hochberg (BH) procedure [Benjamini and Hochberg, 1995] at the nominal level *α* = 0.05.

#### 4.3.3 Conditional sampling via residual permutation

To generate samples from *p*(*X^j^ | X^−j^*), we employed a conditional permutation scheme [Chamma et al., 2023]. For each feature *X^j^*, a conditional model *ν_j_* was trained to predict *X^j^* from *X^−j^* on the training set. For continuous features, *ν_j_* was a ridge regression with regularization strength selected via cross-validation (RidgeCV from scikit-learn). For categorical features, *ν_j_* was an *ℓ*_2_-regularized logistic regression with regularization selected via cross-validation (LogisticRegressionCV, Pedregosa et al. 2011). This choice over more complex models used to predict outcomes is motivated by the fact that relationships between features are easier to predict. Indeed, each feature can be predicted using others collected at the same time point, or with the same modality and are less noisy than outcomes based on cognitive scores. Conditional samples were then generated by permuting the residuals *r_i_* = *X^j^_i_ − ν_j_*(*X^−j^_i_*) across test observations for continuous features, or by sampling the class label from the predicted probability vector *ν_j_*(*X^−j^*) for categorical features. This procedure preserves the estimated conditional mean E[*X^j^ | X^−j^*] while breaking the unique association between *X^j^* and *Y* given *X^−j^*. We used *B* = 10 conditional permutations per feature per cross-validation fold, and averaged the resulting importance estimates to tame variance.

#### 4.3.4 CFI algorithm

The full CFI procedure is summarized in Algorithm 1.

##### Algorithm 1 Conditional Feature Importance (CFI)

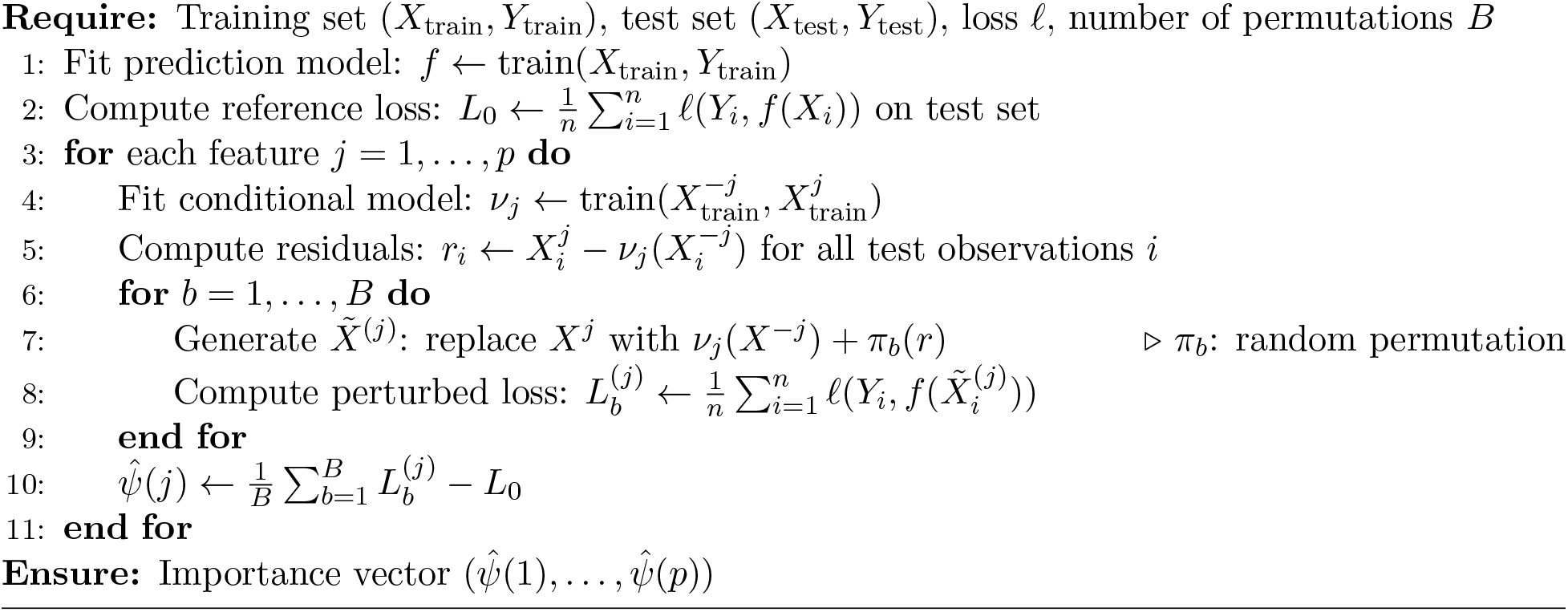

#### 4.3.5 Prediction models

For BioFINDER and ADNI, we used TabICLv2 [Qu et al., 2025] as the primary prediction model, a tabular foundation model based on in-context learning that achieves state-of-the-art performance on tabular classification tasks without task-specific hyperparameter tuning. TabPFNv2.5 [Hollmann et al., 2025] was used as a secondary model to assess robustness of importance estimates across architectures (Figure S4). For the UK Biobank survival analysis, we used CoxNet (elastic-net penalized Cox proportional hazards model) from the scikit-survival library [Pölsterl, 2020], along with univariate screening to retain 100 features with the strongest univariate associations. The screening was part of the cross-validation loop (only using training samples) to avoid data leakage.

#### 4.3.6 Cross-validation and evaluation

All experiments used stratified 10-fold cross-validation, repeated ten times for BioFINDER and ADNI with the same splits across models; UK Biobank used a single 10-fold cross-validation. For BioFINDER classification, stratification was performed jointly on diagnostic group and *APOE ε*4 allele count to ensure balanced representation of this key genetic factor across folds. For CDR-SB regression, stratification used three equal-width intervals of the progression outcome crossed with *APOE ε*4 allele count. For ADNI, stratification was based on diagnostic group. Within each fold, the predicton model *f* and the conditional models 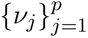 were trained on the training set, and importance was computed on the held-out test set. Final importance scores *ψ*^^^(*j*) were averaged across the 10 folds and, for BioFINDER and ADNI, across the ten repeats. Model performance was evaluated using weighted one-vs-one ROC AUC for multi-class classification, *R*^2^ for regression, and concordance index for survival analysis.

#### 4.3.7 Marginal importance estimation

The marginal importance quantifies the marginal predictive contribution of a feature *X^j^*, including the variance shared with other features but excluding interaction effects. It is estimated using the Leave-One-Covariate In (LOCI) procedure [Reyero-Lobo et al., 2025]. Denoting *S_j_* the marginal importance of a feature *j*, it is computed following,

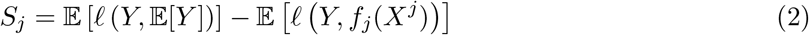

where *f_j_*is a model trained on the *j^th^*feature only and E[*Y* ] corresponds to the mean value for regression tasks and class frequencies for classification tasks.

### 4.4 Feature grouping

#### 4.4.1 Group conditional importance

CFI naturally extends from individual features to groups of features. For a group *G ⊆ {*1*, . . . , p}*, the group conditional importance *ψ*(*G*) is defined as in Equation (1), with *X̃* (*^j^*^)^ replaced by *X̃* (*^G^*^)^, where all features in *G* are jointly sampled from *p*(*X^G^ | X^−G^*). In practice, the conditional model *ν_G_* predicts the entire group *X^G^* from the remaining features *X^−G^*: for groups of continuous features, *ν_G_* is a multivariate ridge regression from *X^−G^*, and their residuals are permuted jointly to preserve within-group correlations. This strategy retains the high correlation within each group, preserving statistical power, while testing whether the group as a whole carries unique predictive information beyond the remaining features. Type-1 error control is maintained at the group level, and BH correction is applied across groups.

#### 4.4.2 Group definitions

We employed three grouping strategies depending on the analysis:

##### Hierarchical clustering

For the dendrogram analysis in BioFINDER (Figure 3 **c**), features were hierarchically clustered using Ward’s method [Ward, 1963] applied to the feature matrix. Group importance was computed at every node of the resulting dendrogram, providing a multi-scale view of which feature clusters carry joint predictive information.

##### Domain-defined groups

For modality-level analyses (Figure 3 **a** and Figure 4 **a**), features were grouped by data source (CSF, plasma, demographics, genetics for BioFINDER; MRI, PET, demographics for ADNI). CSF markers were additionally split into NfL and total tau, which were also measured in plasma, and ten other CSF markers. For pathway-level analysis (Figure 3 **b**), groups were defined as the intersection between published gene sets and the proteins measured in the UK Biobank Olink panel.

##### Data-driven clustering for neuroimaging

For the brain surface maps (Figure 4 **b, c**), cortical regions within each imaging modality (MRI thickness, amyloid PET) were grouped into 15 clusters using Ward’s agglomerative feature clustering (FeatureAgglomeration from scikit-learn), fitted on the standardized feature matrix. This resolves the extreme multicollinearity between neighbouring cortical regions that causes individual-level CFI to lose power. The result of the clustering algorithm on ADNI can be visualized in Figure 5.

## 5 Supplement

**Figure S1:**
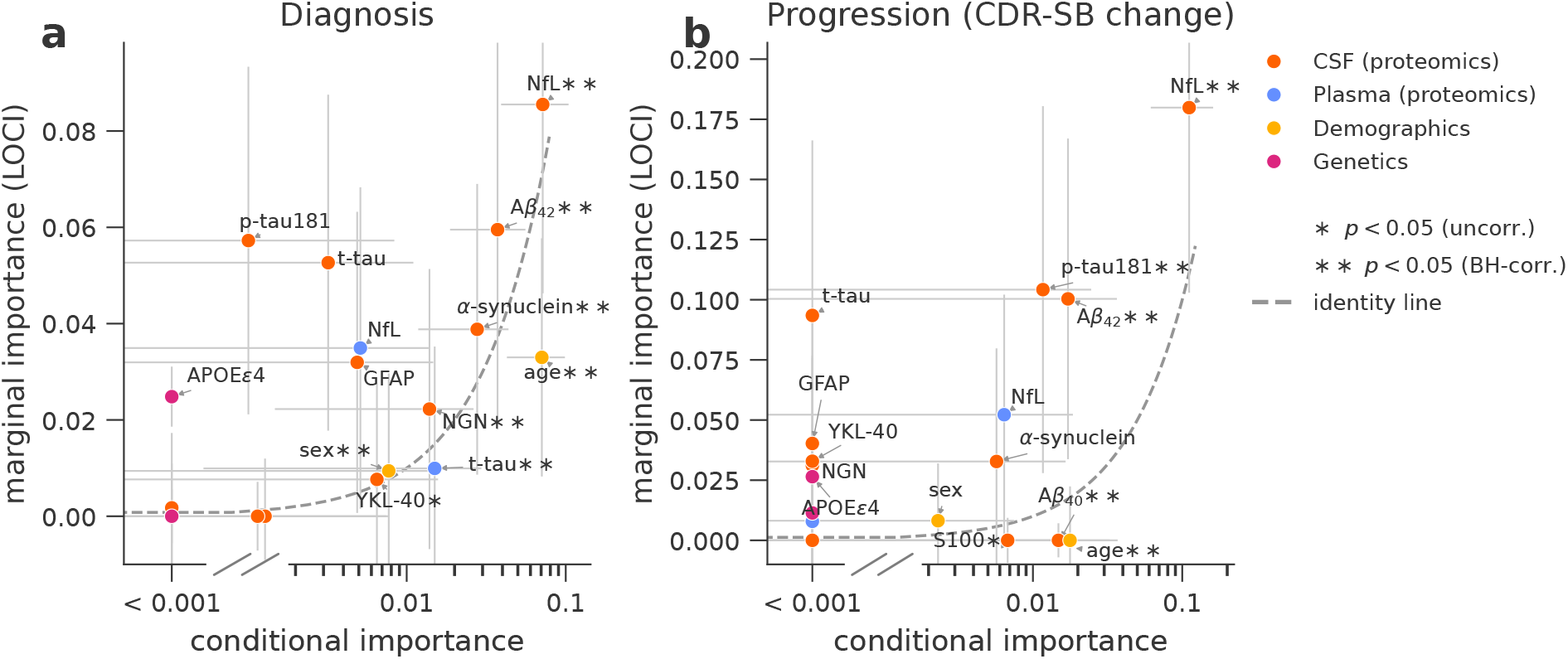
Conditional and marginal feature importance. The plot represents the conditional importance on the horizontal axis and the marginal importance, measured with the Leave One Covariate In (LOCI) approach, on the vertical axis. **(a)** Importances for the AD classification task and **(b)** for the progression prediction task. For each variable, the color encodes the modality (CSF, plasma, demographics, genetics). The error bars represent one standard deviation, estimated across ten repeats of 10-fold cross-validation. Asterisk next to the feature name encodes statistical significance for a one-sided t-test with Nadeau-Bengio correction. A single asterisk corresponds to a *p*-value smaller than 0.05 and two asterisks to a BH-corrected *p*-value below 0.05. Negative conditional importance values were clipped to 0 for visualization only.

**Figure S2:**
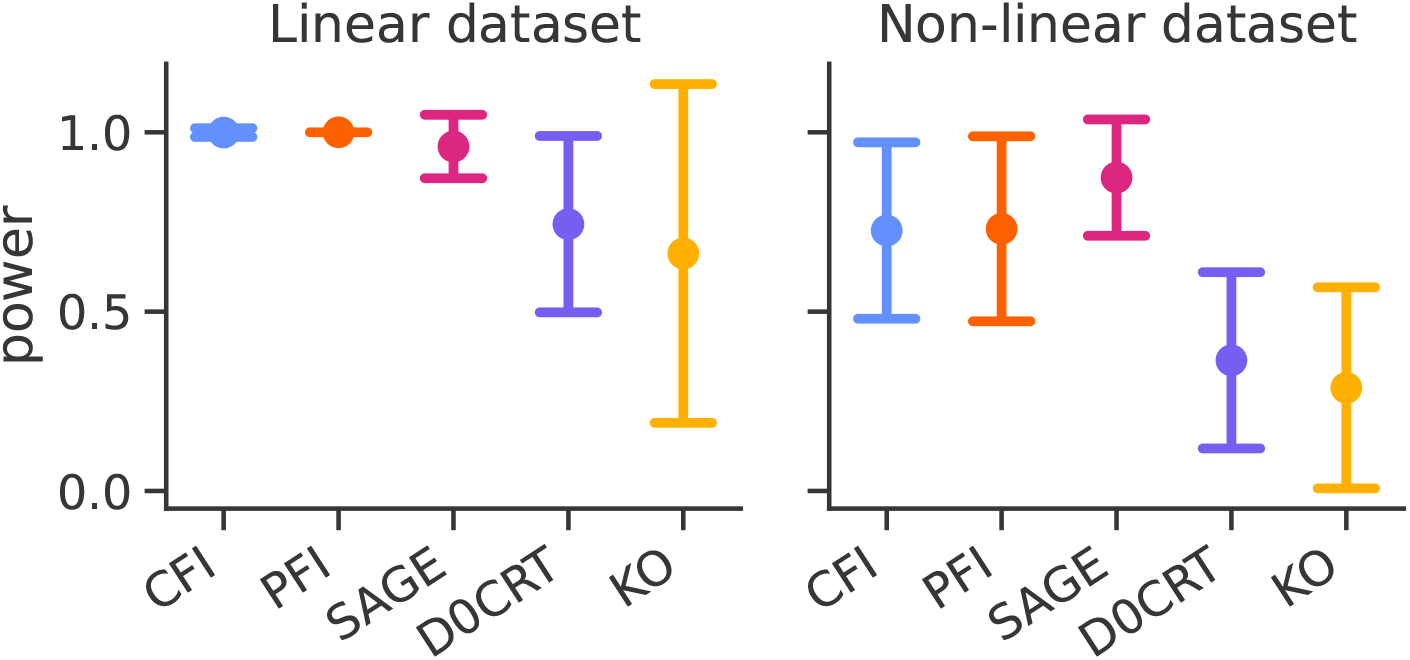
Power comparison across feature importance methods. Shows the distribution of power across 1000 simulations, using the setup from Figure 1. Error bars represent one standard deviation.

**Figure S3:**
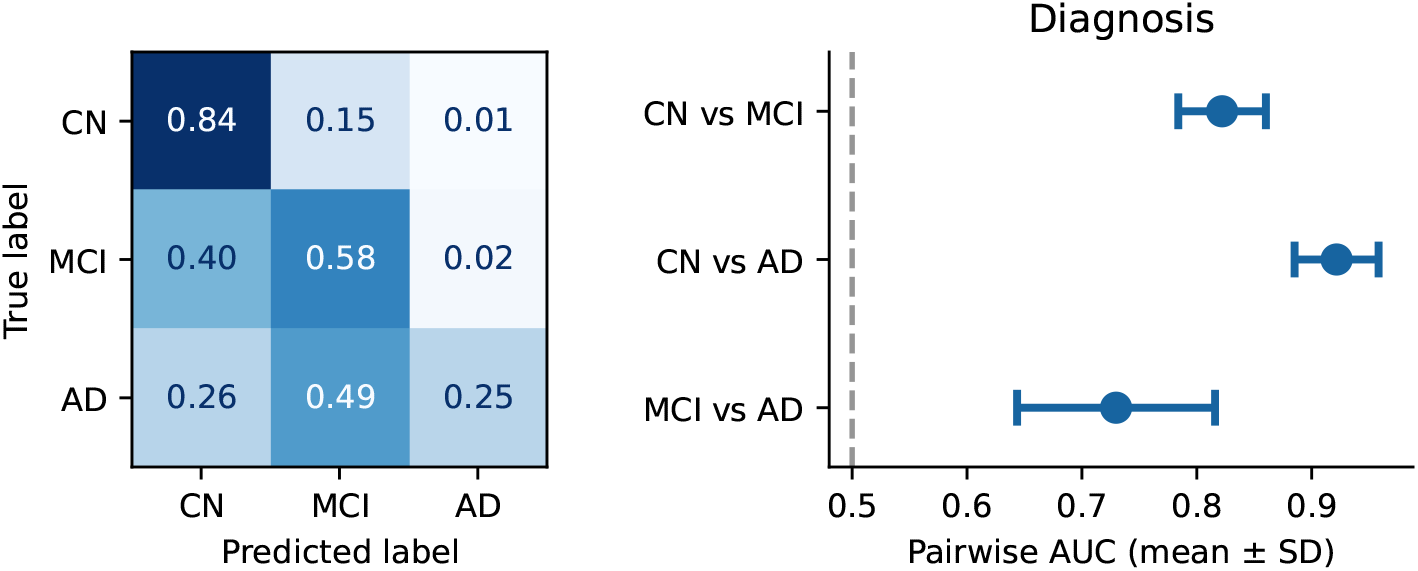
Diagnosis classification on BioFINDER. Provides additional metrics for the classification task using the TabICL model (see Figure 2 (**a**) for the benchmark). (**a**) shows rownormalized prediction rates pooled across ten repeats of 10-fold cross-validation. (**b**) the mean pairwise AUC and one standard deviation across folds for CN versus MCI, CN versus AD, and MCI versus AD. Pairwise AUCs were calculated from the same three-class model by averaging the two class directions.

## 6 Agreement of importance estimates across prediction models

Conditional feature importance was estimated for the 18 BioFINDER variables with three alternative models: a second tabular foundation model (TabPFN), histogram-based gradient boosting (HGB), and a penalized linear model (logistic regression for diagnosis, ridge regression for CDR-SB change). All were fit on the same cross-validation splits with the same conditional sampler as the reference model, TabICLv2, so the comparison is paired. Figure S4 plots the fold-averaged importances of each alternative against the reference, with *R*^2^ the squared Pearson correlation between the two sets of 18 values.

In the diagnosis task, this agreement, measured with *R*^2^, was 0.99 for TabPFN, 0.93 for HGB and 0.74 for logistic regression. In the CDR-SB change task, it was 0.99 for TabPFN, 0.90 for ridge regression and 0.96 for HGB. The cross-validated predictive performance of each model is reported on the corresponding axis of each panel. After BH correction, TabPFN reproduced all seven diagnosis selections made by TabICLv2 and additionally selected plasma NfL; for progression, only CSF NfL was selected by both models.

Conditional feature importance targets the total Sobol index, a property of the data rather than of the model used to estimate it, so models that describe the data well are expected to agree on which variables carry conditional information. Agreement with the reference increased with predictive performance in both tasks, and the looser agreement of the least predictive model reflects conditional signal that this model did not extract.

**Figure S4:**
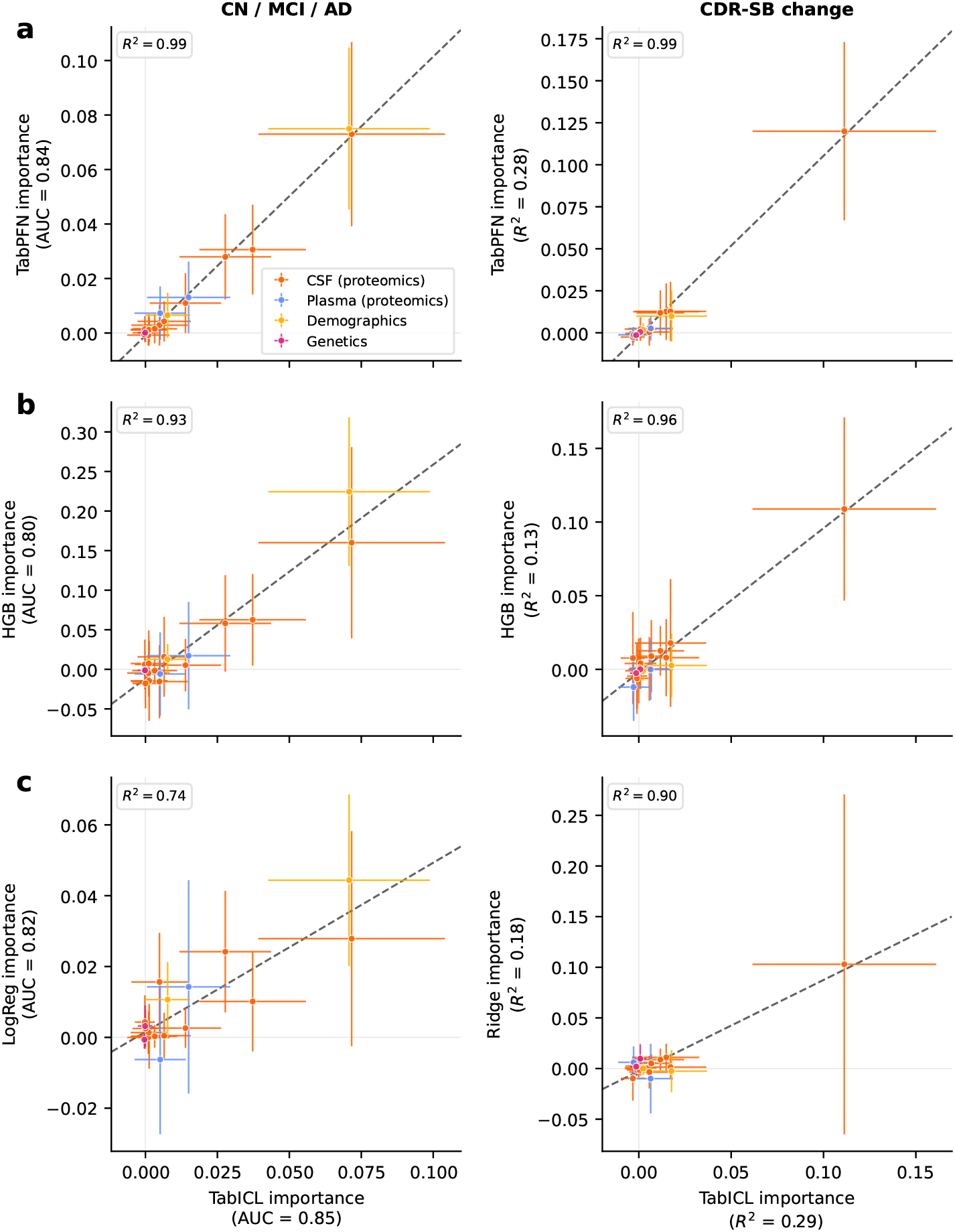
Importance estimates agree across predictive models. Conditional feature importance of the 18 BioFINDER variables estimated with the reference model, TabICL (x-axis), against three alternative models (y-axis): **a**, TabPFN, a second tabular foundation model; **b**, histogram gradient boosting (HGB); and **c**, a penalized linear baseline (logistic regression for diagnosis, ridge regression for CDR-SB change). The left column shows the diagnosis classification task and the right column the CDR-SB change regression task. Each point is one variable, with its importance averaged over ten repeats of 10-fold cross-validation; horizontal and vertical bars show the standard deviation across folds. The dashed grey line is the least-squares linear fit. Colors encode the modality of the variable. We report the coefficient of determination *R*^2^ of that fit, computed over the 18 variables. Axis labels report the mean cross-validated predictive performance of each model on the corresponding task (ROC AUC for diagnosis, *R*^2^ for CDR-SB change). The x-axis is shared across rows.

**Figure S5:**
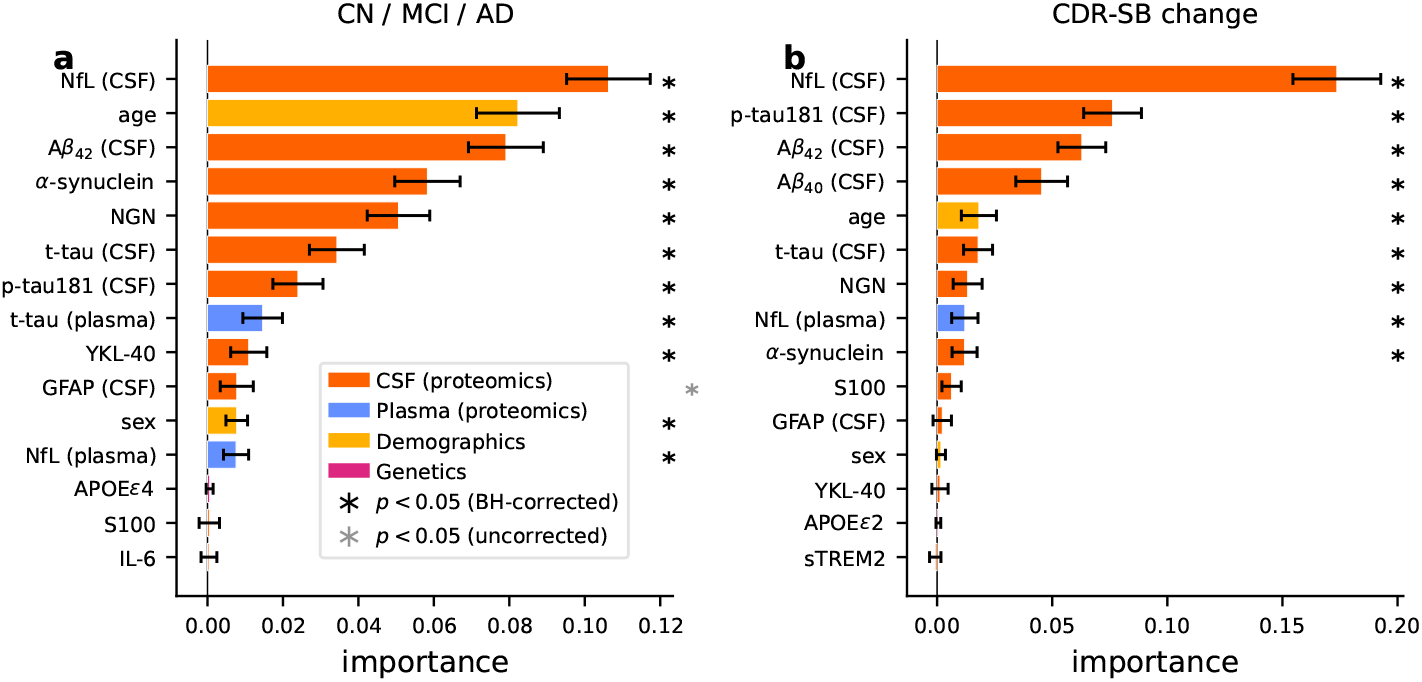
PFI proteomic importances on BioFINDER. (**a**) Bars show mean TabICL permutation importance, with TabPFN estimates shown as comparison markers. Error bars show one standard deviation across 10 times 10 fold cross-validation. Similar to Figure 2 (**b**) but using PFI instead of CFI. (**b**) Individual importances measured with PFI for progression regression on BioFINDER. Similar to Figure 2 (**b**) but using PFI instead of CFI.

**Figure S6:**
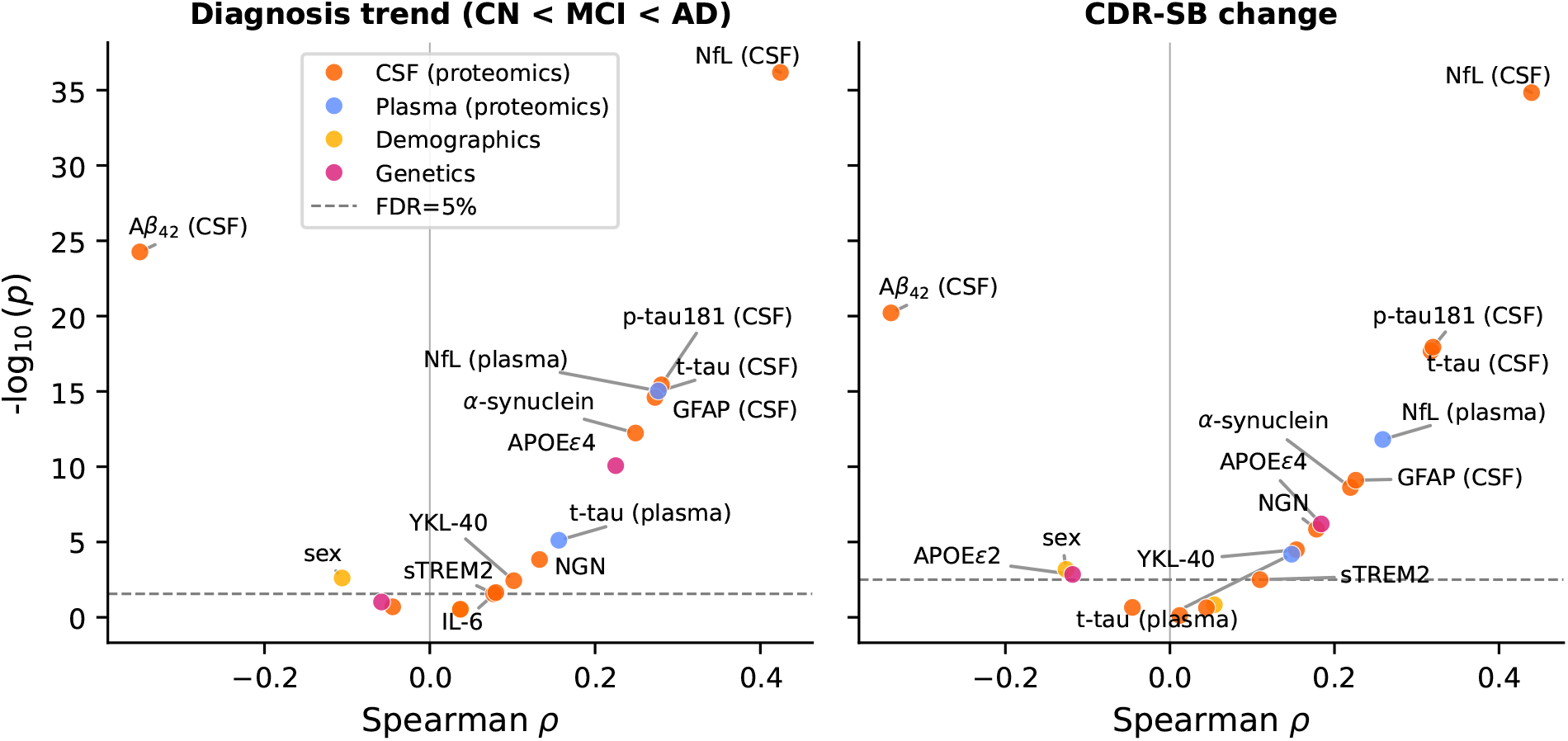
Univariate Spearman correlations. Univariate associations between BioFINDER features and diagnosis (left, coded as CN=0, MCI=1 and AD=2) and rank-normalized CDR-SB progression (right). The x-axis represents the Spearman correlation, and the y-axis *−* log_10_ of the corresponding p-value. The horizontal dashed gray line represents the BH threshold at the level *α* = 0.05, applied separately to each outcome. The colors encode the different modalities.

## 7 Composition of the pathway groups in the UK Biobank

Table S1 lists the composition of the pathway groups used in Figure 3 **b**, each group being the intersection between a gene set [Ekins et al., 2007] and the proteins measured by the Olink panel.

**Table S1:**
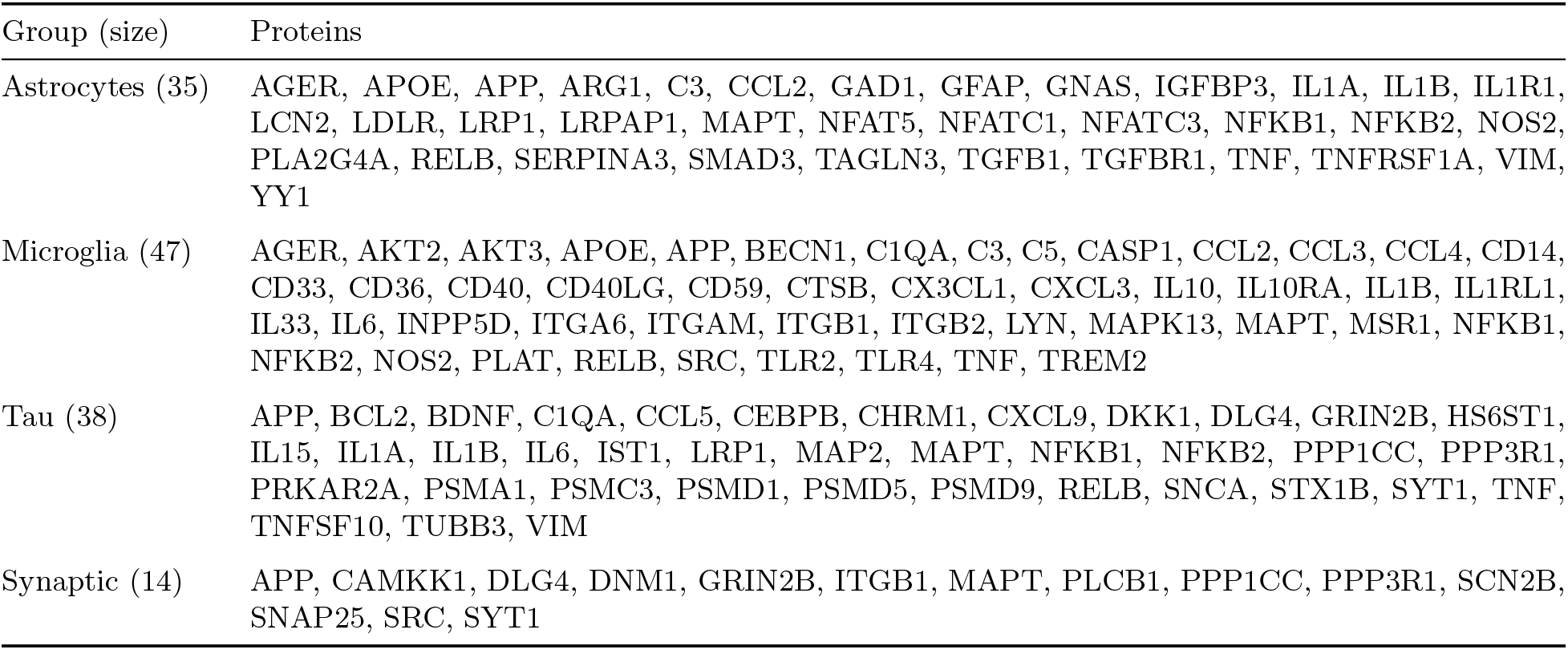
Composition of the pathway groups used for group-level conditional feature importance in the UK Biobank (Figure 3 b). Each group is the intersection between a gene set and the proteins measured by the Olink panel; only the intersection is listed, and group-level CFI was computed on these proteins. Proteins are named by gene symbol, as in Figure 2 **e**; APOE denotes the plasma protein rather than the *APOE ε*4 genotype. The demographics group (age and sex) is not shown.

| Group (size) | Proteins |
|---|---|
| Astrocytes (35) | AGER, APOE, APP, ARG1, C3, CCL2, GAD1, GFAP, GNAS, IGFBP3, IL1A, IL1B, IL1R1, LCN2, LDLR, LRP1, LRPAP1, MAPT, NFAT5, NFATC1, NFATC3, NFKB1, NFKB2, NOS2, PLA2G4A, RELB, SERPINA3, SMAD3, TAGLN3, TGFB1, TGFBR1, TNF, TNFRSF1A, VIM, YY1 |
| Microglia (47) | AGER, AKT2, AKT3, APOE, APP, BECN1, C1QA, C3, C5, CASP1, CCL2, CCL3, CCL4, CD14, CD33, CD36, CD40, CD40LG, CD59, CTSB, CX3CL1, CXCL3, IL10, IL10RA, IL1B, IL1RL1, IL33, IL6, INPP5D, ITGA6, ITGAM, ITGB1, ITGB2, LYN, MAPK13, MAPT, MSR1, NFKB1, NFKB2, NOS2, PLAT, RELB, SRC, TLR2, TLR4, TNF, TREM2 |
| Tau (38) | APP, BCL2, BDNF, C1QA, CCL5, CEBPB, CHRM1, CXCL9, DKK1, DLG4, GRIN2B, HS6ST1, IL15, IL1A, IL1B, IL6, IST1, LRP1, MAP2, MAPT, NFKB1, NFKB2, PPP1CC, PPP3R1, PRKAR2A, PSMA1, PSMC3, PSMD1, PSMD5, PSMD9, RELB, SNCA, STX1B, SYT1, TNF, TNFSF10, TUBB3, VIM |
| Synaptic (14) | APP, CAMKK1, DLG4, DNM1, GRIN2B, ITGB1, MAPT, PLCB1, PPP1CC, PPP3R1, SCN2B, SNAP25, SRC, SYT1 |

## Footnotes

1 https://github.com/mind-inria/hidimstat

